## Supplementary Information for "Illuminating the mechanism and allosteric behavior of NanoLuc luciferase"

1. Loschmidt Laboratories, Department of Experimental Biology and RECETOX, Faculty of Science, Masaryk University, Kamenice 5, Bld. C13, 625 00 Brno, Czech Republic
2. International Clinical Research Center, St. Anne's University Hospital Brno, Pekarska 53, 656 91 Brno, Czech Republic
3. Current address: Center for Interdisciplinary Biosciences, Technology and Innovation Park, P. J. Safarik University in Kosice, Trieda SNP 1, 04011 Kosice, Slovakia
4. Department of Histology and Embryology. Faculty of Medicine, Masaryk University, Kamenice 753/5, 625 00 Brno, Czech Republic
5. Unité de Chimie et Biocatalyse, Institut Pasteur, UMR 3523, CNRS, 28 rue du Dr. Roux, 75724 Paris Cedex 15, France
6. Université de Paris, 12 rue de l'école de Médecine, 75006 Paris, France.
7. Structure et Instabilité des Génomes (StrInG), Muséum National d'Histoire Naturelle, INSERM, CNRS, Alliance Sorbonne Université, 75005 Paris, France

[#] These authors contributed equally to this study.

### Contents

Supplementary figures and tables ………………………………………………………………. Pages 2 to 30

Supplementary schemes ………………………………………………………….……………...…. Page 32

Supplementary notes ……………………………………………………………………..………. Pages 33 to 34

**Supplementary Table 1.** Crystallographic data collection and refinement statistics.

| | NanoLuc/FMA | NanoLuc/CEI | NanoLuc-Y94A | NanoLuc$^{CTZ}$/azaCTZ |
|---|---|---|---|---|
| **Data collection** | | | | |
| Wavelength (Å) | 1 | 1 | 1 | 1 |
| Space group | *C*121 | *C*121 | *C*222 | *C*121 |
| Cell dimensions | | | | |
| a, b, c (Å) | 86.91, 87.49, 191.76 | 86.98, 87.28, 191.52 | 86.45, 86.47, 96.24 | 147.91, 112.6, 59.7 |
| α, β, γ (°) | 90, 90.09, 90 | 90, 90.06, 90 | 90, 90, 90 | 90, 90.8, 90 |
| Resolution (Å) | 47.94 - 1.70 (1.73 - 1.70) | 47.88 - 2.0 (2.03 - 2.0) | 48.12 - 2.8 (2.95 - 2.8) | 49.49 - 3.1 (3.31 - 3.1) |
| Total reflections | 528,509 (24,117) | 312,703 (12,347) | 59,773 (8,927) | 61,192 (11,596) |
| Unique reflections | 158,483 (14,059) | 96,944 (8,970) | 9,192 (897) | 17,550 (3,209) |
| Rmerge | 0.066 (0.786) | 0.087 (0.839) | 0.152 (1.697) | 0.107 (2.487) |
| I/σI | 9.7 (1.5) | 8.7 (1.2) | 10.6 (1.3) | 6 (0.7) |
| Completeness (%) | 99.3 (94.5) | 98.9 (92.5) | 99.9 (99.9) | 98.5 (99.5) |
| Multiplicity | 3.4 (3.3) | 3.3 (2.8) | 6.5 (6.9) | 3.5 (3.6) |
| CC(1/2) | 0.996 (0.503) | 0.997 (0.36) | 0.997 (0.28) | 0.997 (0.22) |
| Wilson B-factor | 25.0 | 32.1 | 74.9 | 111.3 |
| **Refinement** | | | | |
| Resolution (Å) | 47.94 - 1.69 (1.751 - 1.69) | 47.88 - 2.0 (2.03 - 2.0) | 48.12 - 2.8 (2.95 - 2.8) | 49.49 - 3.1 (3.31 - 3.1) |
| No. reflections | 158,483 (14,059) | 96,944 (8,970) | 9,192 (897) | 15,549 (1,540) |
| No. reflections for R-free | 8,057 (682) | 4,974 (424) | 430 (41) | 894 (75) |
| Rwork / Rfree (%) | 13.45 / 18.27 | 18.84 / 20.74 | 26.14 / 30.48 | 27.24 / 30.74 |
| No. atoms | | | | |
| Protein | 10,826 | 10,835 | 2,702 | 4,101 |
| Ligand | 165 | 116 | 0 | 96 |
| Water | 771 | 340 | 16 | 20 |
| B-factors | 33.93 | 37.13 | 67.28 | 109.63 |
| Protein | 33.44 | 37.34 | 67.44 | 109.81 |
| Ligand | 40.69 | 23.07 | | 104.32 |
| Water | 39.44 | 35.48 | 40.75 | 98.5 |
| R.m.s. deviations | | | | |
| Bond lengths (Å) | 0.019 | 0.019 | 0.003 | 0.003 |
| Bond angles (°) | 2.32 | 2.47 | 0.68 | 0.68 |
| Ramachandran favored (%) | 94.16 | 89.89 | 84.91 | 89.47 |
| Ramachandran allowed (%) | 4.58 | 8.04 | 11.83 | 7.21 |
| Ramachandran outliers (%) | 1.26 | 2.07 | 3.25 | 3.31 |
| PDB ID code | 8AQ6 | 8AQI | 8AQH | 8BO9 |

Values in parentheses are for the highest-resolution shell.

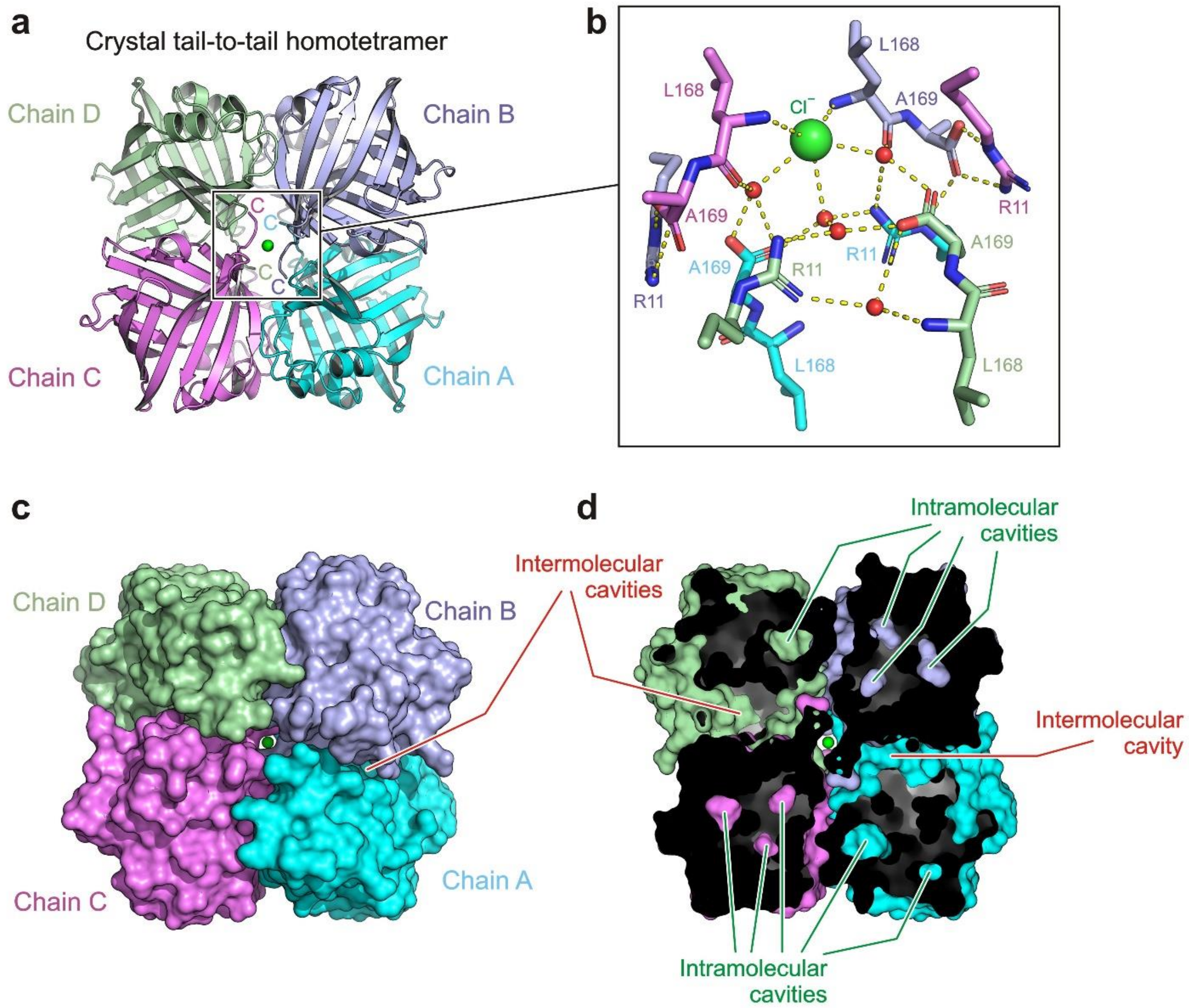


**Supplementary Fig. 1.** Anatomy of NanoLuc crystals. (a) Cartoon representation of crystallographic homotetramer of NanoLuc enzyme with a central pore occupied by a chloride ion (green sphere). Note that all four C-terminal ends are involved in the tight packing. (b) Close-up view of interactions between all four C-terminal ends, chloride ion (green sphere) and solvent (red spheres). (c) Surface representation of crystallographic homotetramer of NanoLuc enzyme with a central pore occupied by a chloride ion (green sphere). Note the tight association of NanoLuc enzyme (chain A to D) in the crystallographic tetramer. (d) Cutaway surface representation of crystallographic NanoLuc homotetrameric association. Note that apart from the intramolecular cavities, there are numerous intermolecular surface pocket.

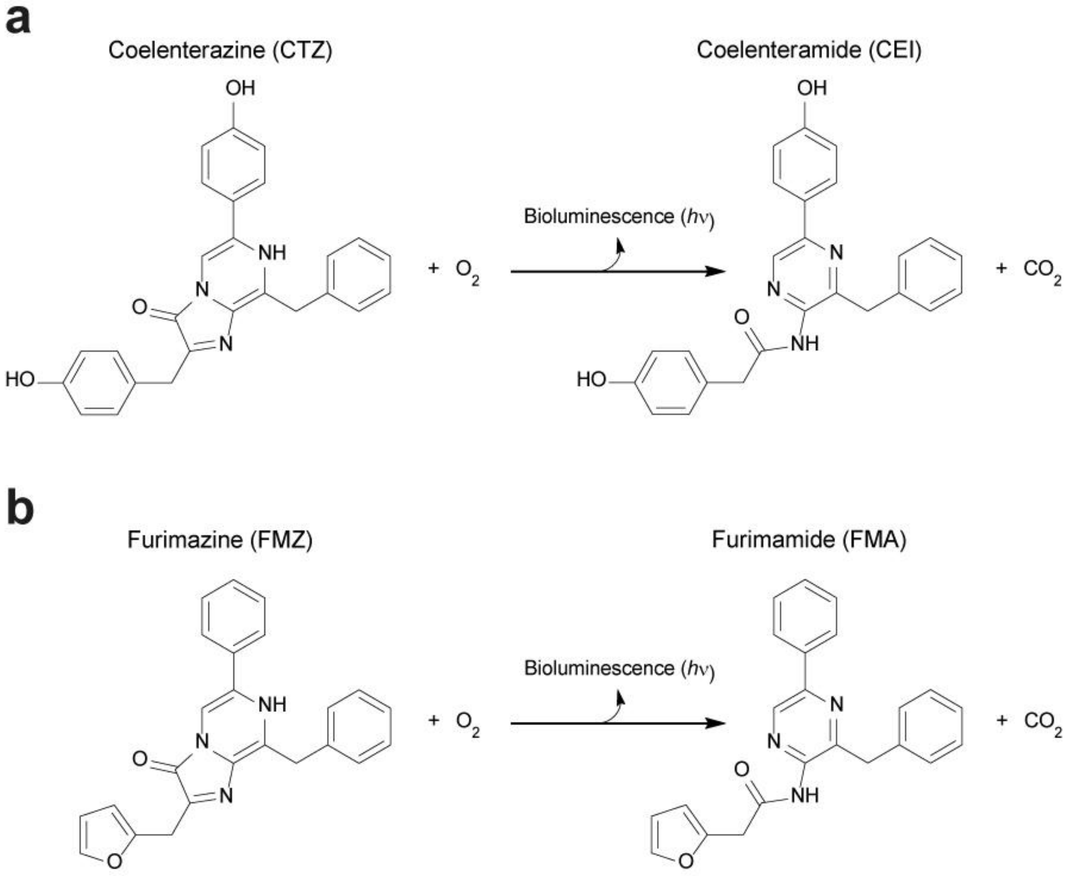


**Supplementary Fig. 2.** Schematic representation of bioluminescence reaction catalyzed by luciferase. The chemical structures of the luciferase substrates coelenterazine (a) and furimazine (b) with their oxidized products coelenteramide and furimamide are indicated.

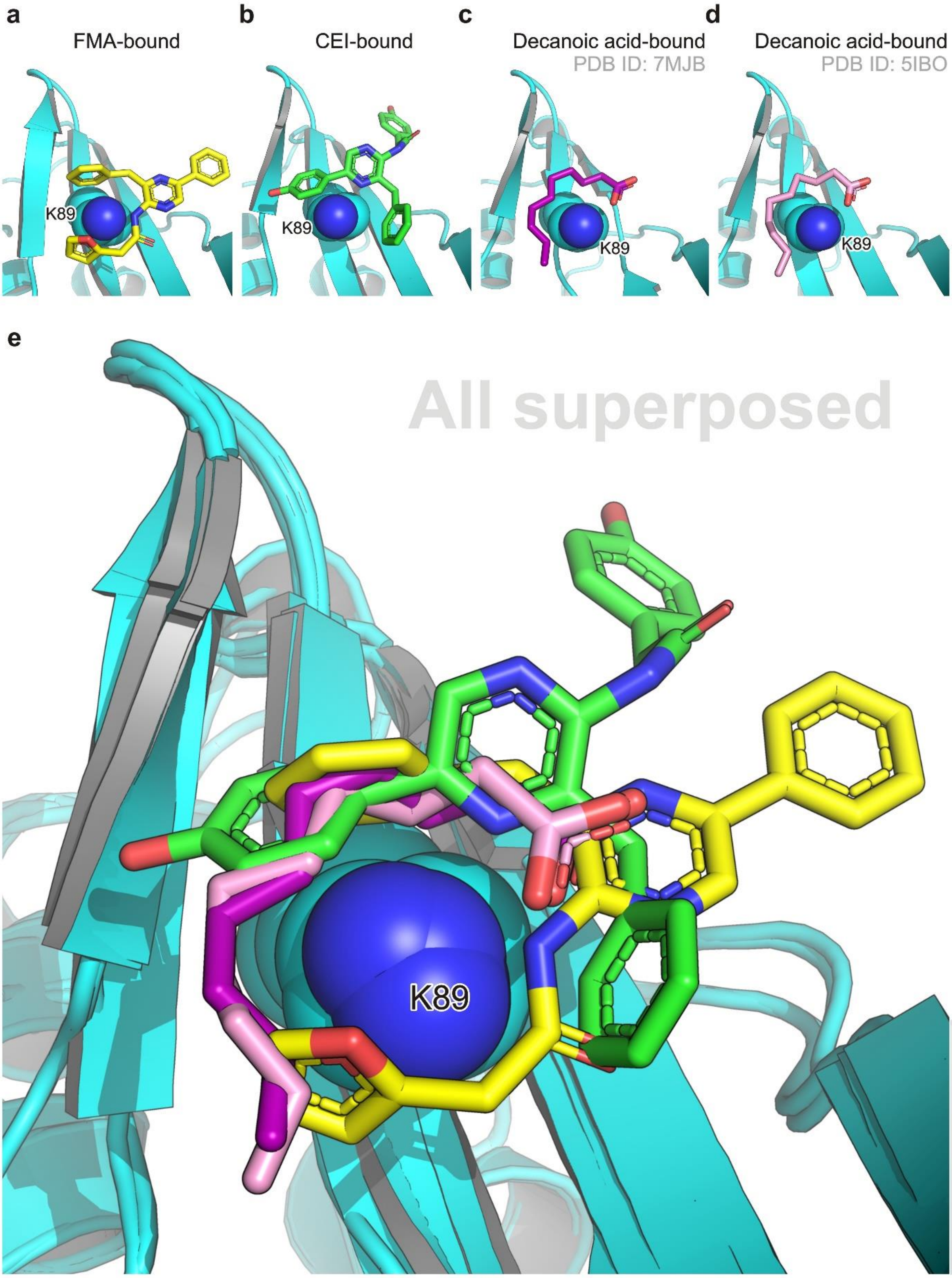


**Supplementary Fig. 3.** Structural comparison of FMA, CEI and decanoic acid binding modes when bound to the NanoLuc surface pocket. (a) Snapshot of FMA (yellow sticks) bound to NanoLuc. (b) Snapshot of CEI (green sticks) bound to NanoLuc. (c) Snapshot of decanoic acid (purple sticks) bound to NanoLuc R162Q mutant (PDB ID: 7MJB). (d) Snapshot of decanoic acid (violet sticks) bound to NanoLuc (PDB ID: 5IBO). (e) Superposition of all complex structures (A to D). In all panels, lysine 89 (K89) is shown in space-filling sphere representation.

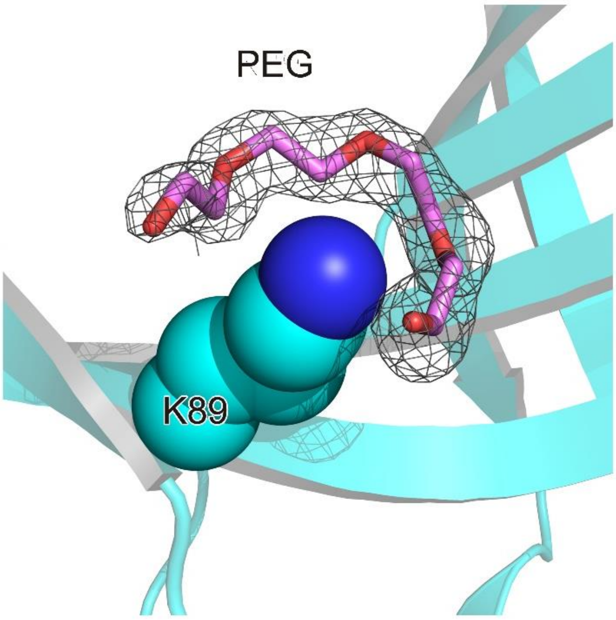


**Supplementary Fig. 4.** Binding of a polyethylene glycol (PEG) molecule at the surface ligand-binding pocket. 2Fo-Fc electron density (contour level 1.2 σ) of PEG molecule (violet sticks), the side chain of K89 is shown as space-filling spheres.

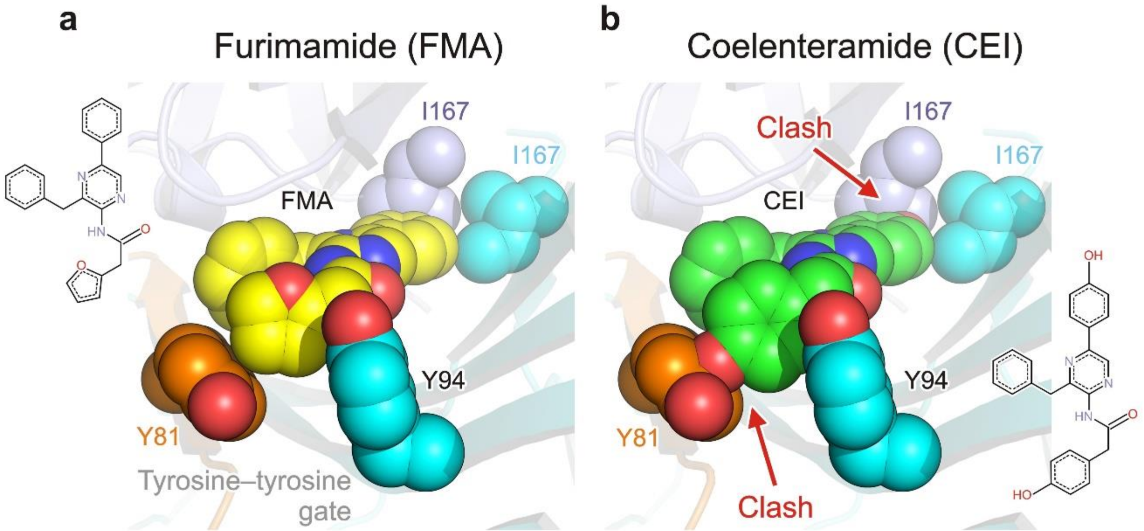


**Supplementary Fig. 5.** Molecular specificities distinguishing FMA-luciferin from CEI-luciferin in the context of the luciferin-binding surface pocket. (a) Close-up view of FMA (yellow) when bound in the NanoLuc surface pocket. Note that the FMA 2-(furan-2-yl) substituent is locked by a tyrosine-tyrosine gate formed by the side chains of Y81 and Y94 residues, while the 8-benzyl substituent is complementary with the hydrophobic environment (I167) of the pocket bottom (b) Close-up view of the modeled CEI (green) molecule in the FMA-preferred binding mode. Note that the CEI 2-(*p*-hydroxyphenyl)acetamide moiety would clash with the side chain of Y81, while the terminal 5-*p*-hydroxyphenyl group would be in a clash with I167 residue in the pocket bottom.

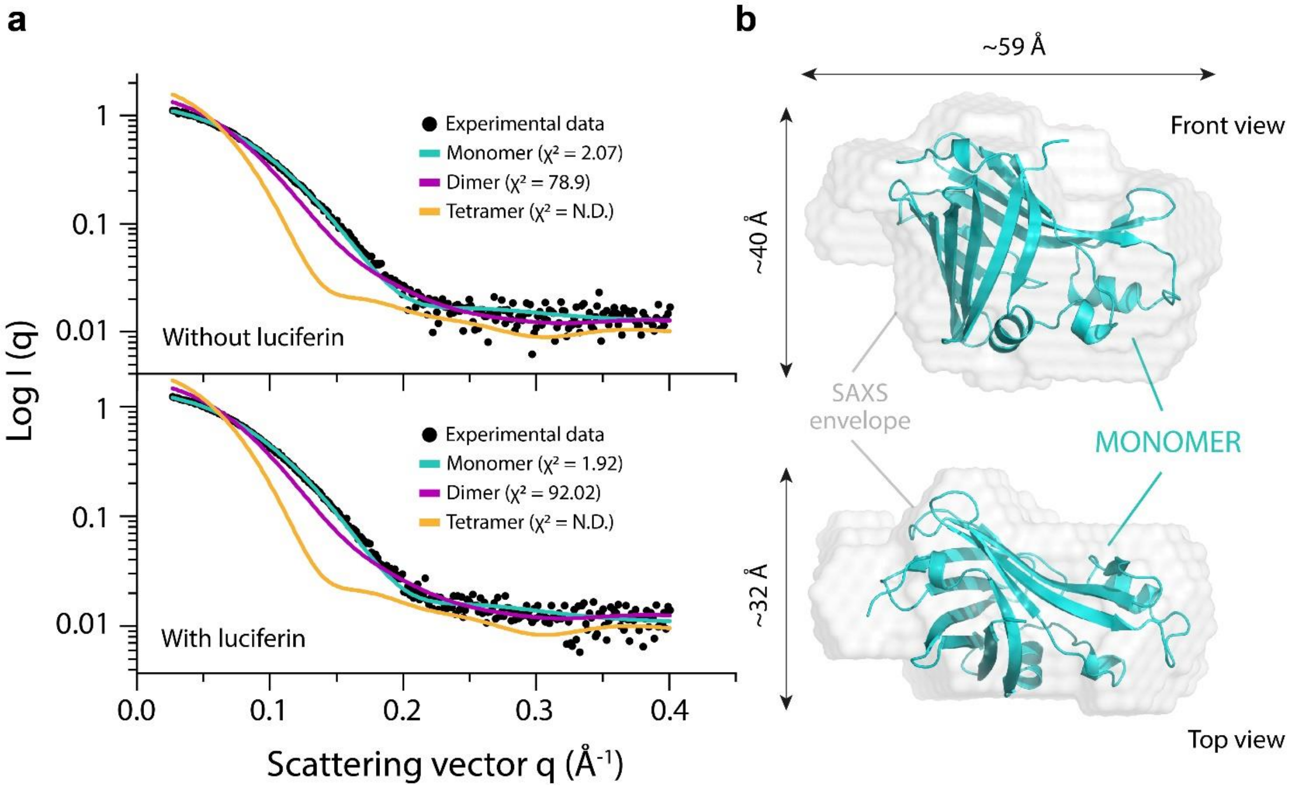


**Supplementary Fig. 6.** Solution structure of NanoLuc determined by SAXS. (a) Experimental SAXS scattering curve for NanoLuc (black dots) is shown against the calculated scattering curves for the NanoLuc monomer (cyan line), dimer (violet line), and tetramer (yellow line). The SAXS curves collected in absence of luciferin (top panel) and in presence of 4 molar excess of FMZ-luciferin (bottom panel) are shown. (b) *Ab initio* molecular envelope generated from SAXS data analysis. The molecular SAXS envelope of the NanoLuc monomer is shown in a semi-transparent grey color superposed on the NanoLuc monomer of the crystal structure represented as a cyan cartoon.

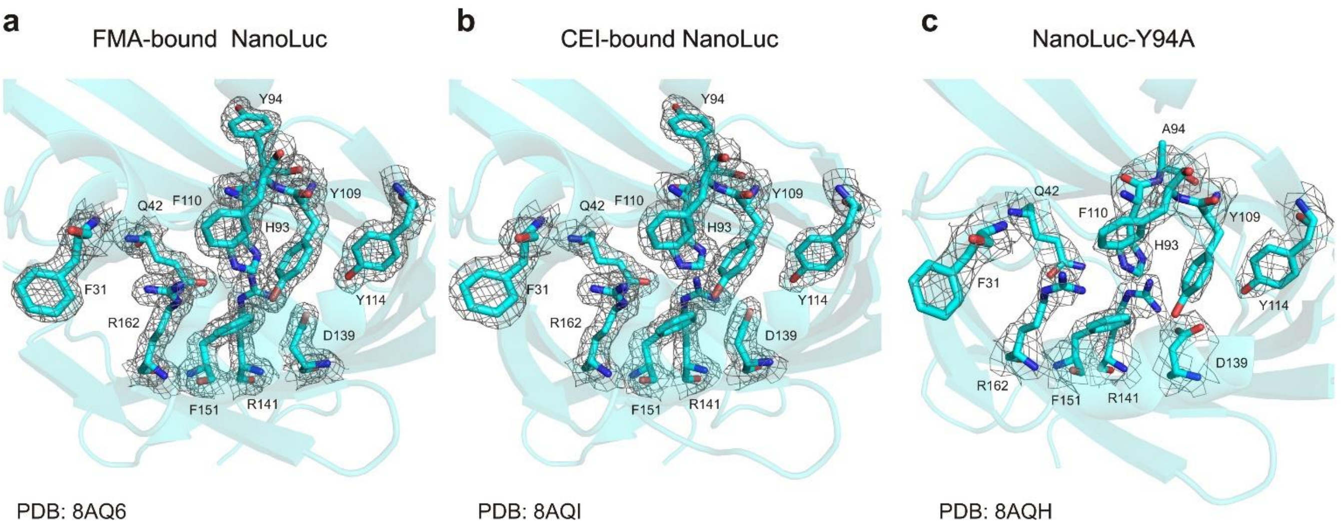


**Supplementary Fig. 7.** 2Fo-Fc electron density maps (contour level 3.0 σ) of catalytic central cavity of FMA-bound (a), CEI-bound (b) and Y94A mutant (c) NanoLuc crystal structures. Note that all structures represent the so-called closed β-barrel state with eliminated central cavity preventing the luciferin binding inside.

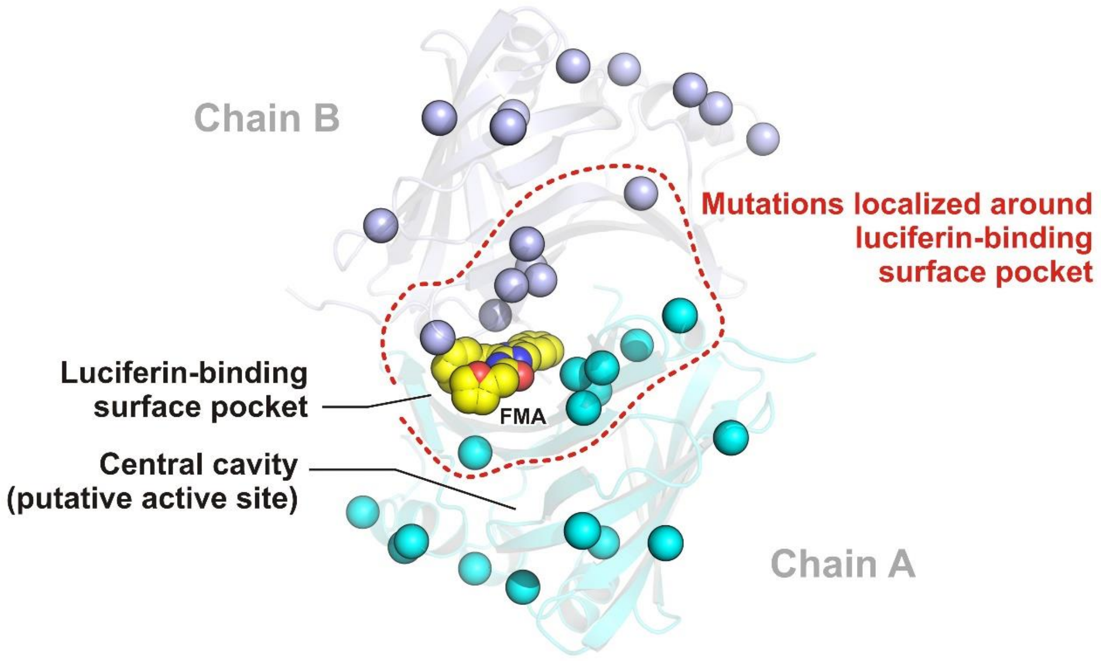


**Supplementary Fig. 8.** Structural distribution of 16 mutated residues during the engineering of NanoLuc luciferase. The FMA-bound asymmetric NanoLuc is shown as cartoon model, chain A as cyan cartoon and chain B as blue cartoon. The FMA luciferin is shown as yellow space-filling spheres; the mutated residues are shown as cyan and blue spheres, respectively.

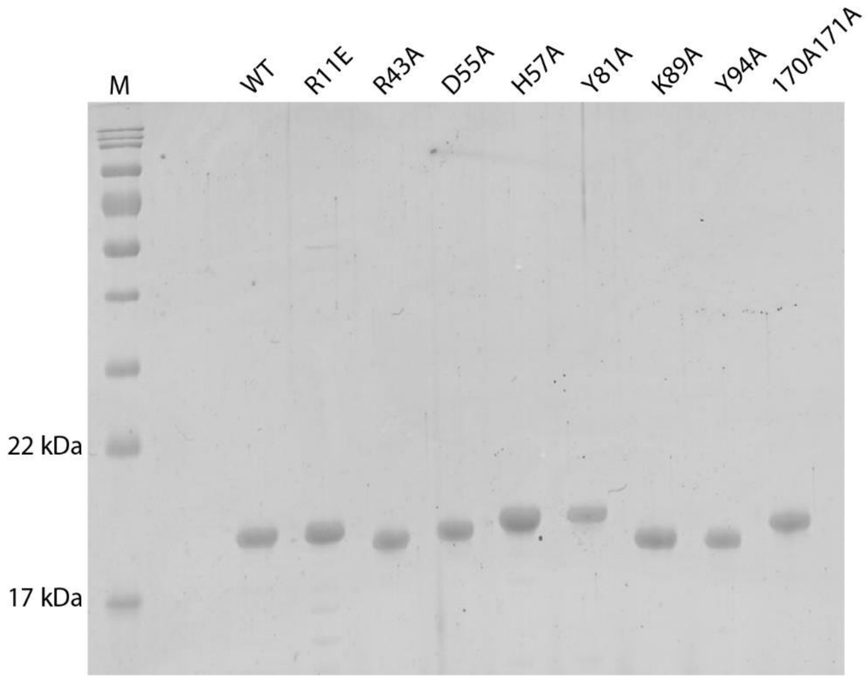


**Supplementary Fig. 9.** SDS-PAGE analysis of NanoLuc variants showing that the constructed mutants were expressed and purified as soluble proteins.

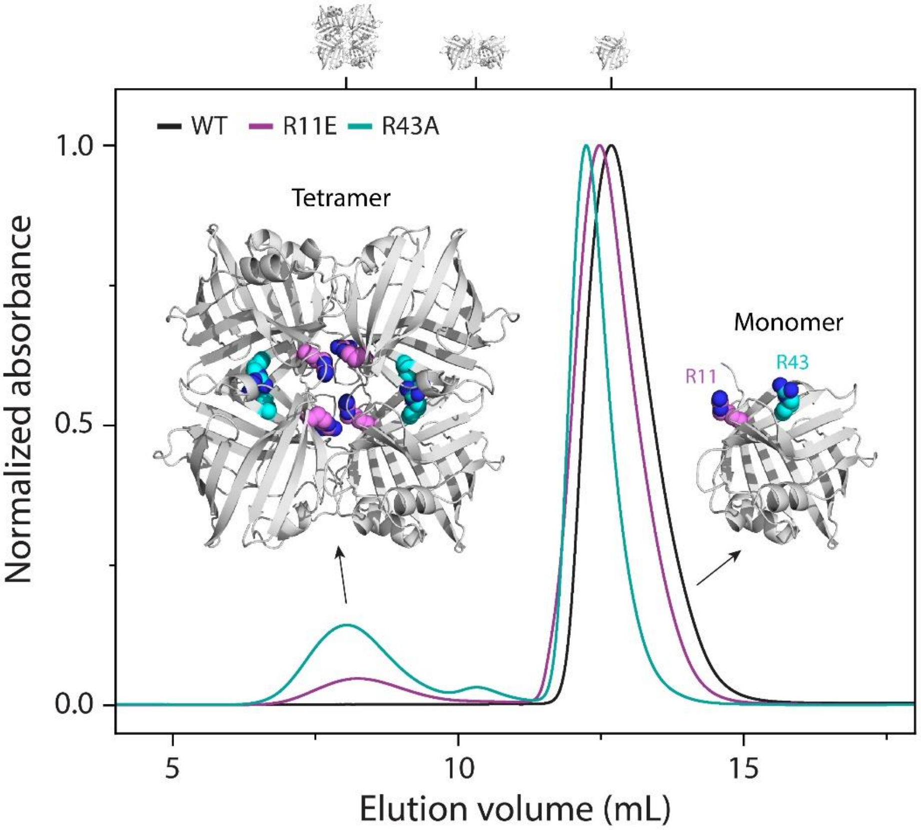


**Supplementary Fig. 10.** Gel filtration purifications of wild-type NanoLuc, NanoLuc-R11E and NanoLuc-R43A. Note that NanoLuc-R11E and NanoLuc-R43A did not exist as a pure monomeric protein anymore but rather as monomer-tetramer mixtures. The mutations are localized at the protein-protein interfaces as depicted in associated structural models.

**Supplementary Table 2.** Luciferase activities of NanoLuc and its mutants with CTZ and FMZ luciferins. Data indicate the average of relative luciferase activity (NanoLuc wild-type = 100%), the error of measurement is represented by the standard deviation (SD).

| NanoLuc variant | CTZ-luminescence | | FMZ-luminescence | |
|---|---|---|---|---|
| | Relative activity (%) | SD (%) | Relative activity (%) | SD (%) |
| Wild-type | 100 | 27.6 | 100 | 2.9 |
| D9R/K89R | 1100.1 | 189.2 | 86.8 | 13.1 |
| D9R/H57A/K89R | 859.8 | 210.1 | 87.3 | 16.3 |
| H57A/K89R | 524.6 | 124.1 | 135.3 | 28.9 |
| H57A | 478.5 | 262.4 | 113.5 | 45.5 |
| K89R | 419.3 | 143.4 | 109.7 | 29.4 |
| D9R/H57A | 346.0 | 49.3 | 28.1 | 5.3 |
| D9R | 272.1 | 112.5 | 52.5 | 24.5 |
| R11E | 138.5 | 1.9 | 97.1 | 2.9 |
| D55A | 109.3 | 26.1 | 56.4 | 2.1 |
| R166A | 104.5 | 50.0 | 91.3 | 33.1 |
| K89E | 97.9 | 6.8 | 44.1 | 3.1 |
| R43A | 76.1 | 26.3 | 59.6 | 15.5 |
| H57Y | 65.9 | 13.3 | 63.5 | 7.1 |
| K89A | 36.3 | 13.8 | 61.2 | 5.0 |
| Y81A | 35.7 | 7.2 | 73.5 | 4.7 |
| R166Q | 22.9 | 9.6 | 73.5 | 0.1 |
| del I167-A169 | 20.6 | 4.6 | 70.1 | 4.4 |
| ext A170-A171 | 22.0 | 1.8 | 43.2 | 2.5 |
| Y94A | 2.0 | 0.4 | 4.6 | 2.2 |

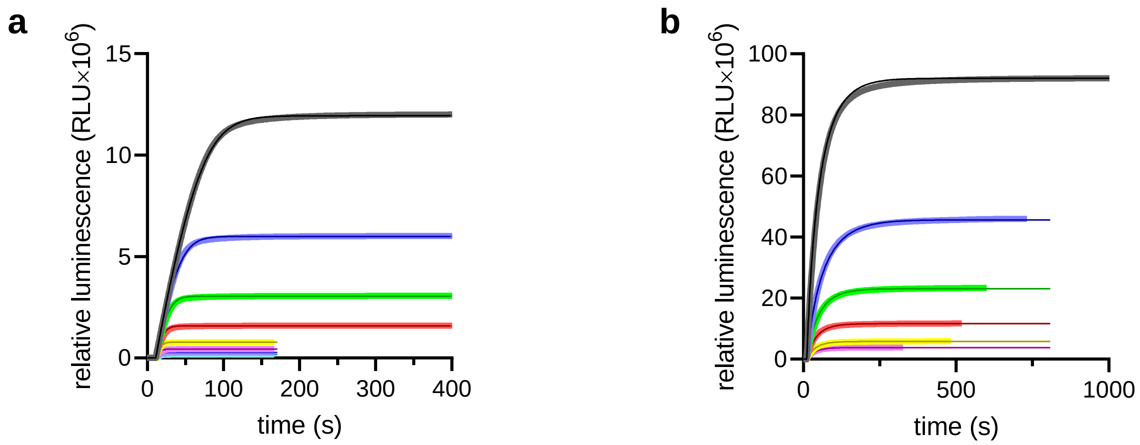


**Supplementary Fig. 11.** Numerical analysis of kinetic parameters of NanoLuc catalyzed luciferin conversion. (a) The reaction progress curves corresponding to cumulative luminescence production in time recorded upon mixing 0.01 µM NanoLuc with 3.88 (black), 1.95 (dark blue), 0.99 (green), 0.51 (red), 0.25 (yellow), 0.14 (magenta), 0.09 (blue) and 0.06 (cyan) µM FMZ, with gain of the reader set to 1000. (b) The reaction progress curves obtained upon mixing 0.02 µM NanoLuc with 0.98 µM CTZ (black) and 0.01 µM NanoLuc with 0.49 (dark blue), 0.25 (green), 0.13 (red) , 0.06 (yellow) and 0.04 (magenta) µM CTZ, with gain of the reader set to 1500. Each trace represents an average of three repetitions. The thicker lines represent the experimental data, thinner lines represent the best fit using reaction model in Scheme 1(see Methods section Measurement of steady-state kinetic parameters of luciferase reaction).

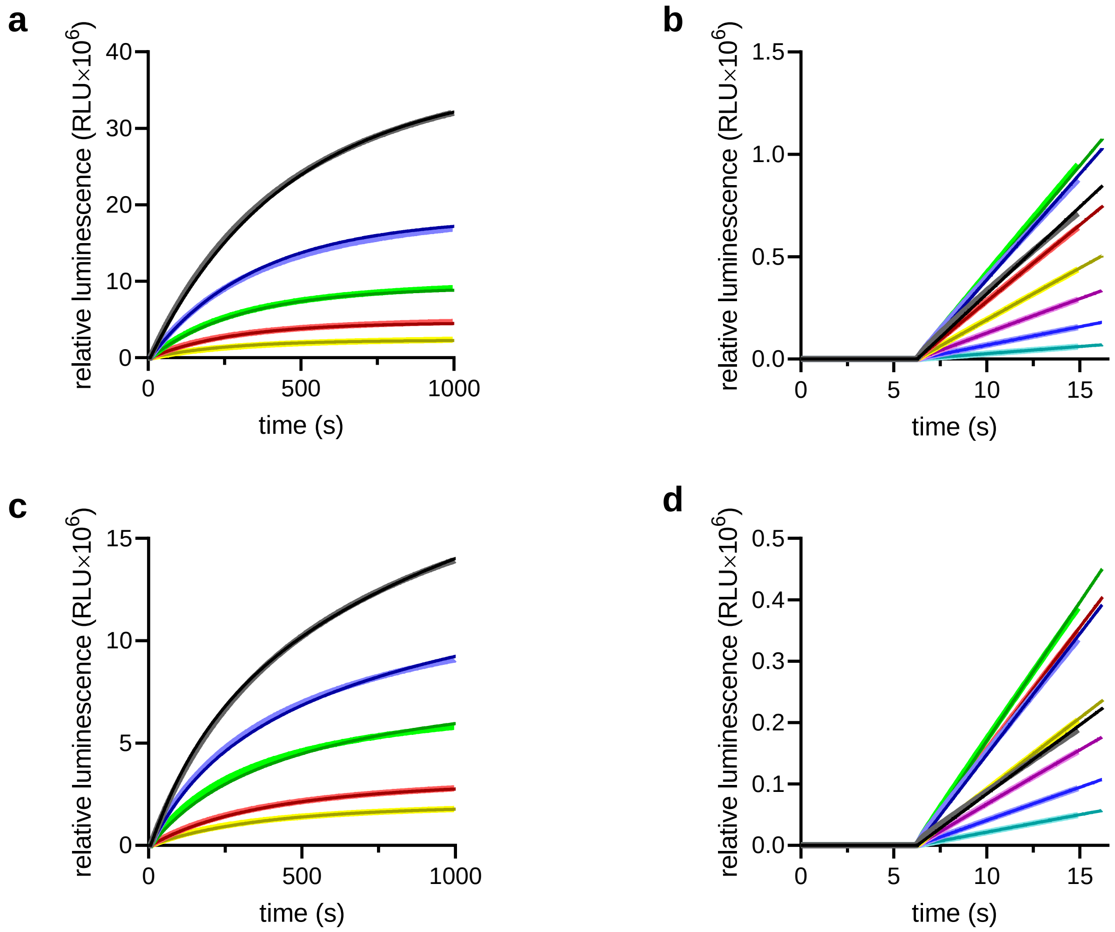


**Supplementary Fig. 12.** Numerical analysis of kinetic parameters of NanoLuc-Y94A catalyzed luciferin conversion. (a) The reaction progress curves corresponding to cumulative luminescence production in time recorded upon mixing 0.01 µM NanoLuc-Y94A with 0.90 (black), 0.45 (dark blue), 0.23 (green), 0.11 (red) and 0.05 (yellow) µM FMZ, with gain of the reader set to 2000. (b) The reaction progress curves capturing the initial reaction velocity of the luciferase reaction corresponding to cumulative luminescence production in time recorded upon mixing 0.01 µM NanoLuc Y94A with 7.18 (black), 3.61 (dark blue), 1.81 (green), 0.89 (red), 0.45 (yellow), 0.23 (magenta), 0.11 (blue) and 0.06 (cyan) µM FMZ, with gain of the reader set to 2000. (c) The reaction progress curves obtained upon mixing 0.05 µM NanoLuc-Y94A with 2.59 (black), 1.66 (dark blue), 0.87 (green), 0.35 (red) and 0.21 (yellow) µM CTZ, with gain of the reader set to 2300. (d) The reaction progress curves capturing the initial reaction velocity of the luciferase reaction corresponding to cumulative luminescence production in time recorded upon mixing 0.05 µM NanoLuc-Y94A with 28.82 (black), 14.45 (dark blue), 7.23 (green), 3.67 (red), 1.81 (yellow), 0.90 (magenta), 0.45 (blue) and 0.23 (cyan) µM coelenterazine, with gain of the reader set to 2300. Each trace represents an average of three repetitions. The thicker lines represent the experimental data, thinner lines represent the best fit using reaction model in Scheme 2 (see Methods section Measurement of steady-state kinetic parameters of luciferase reaction).

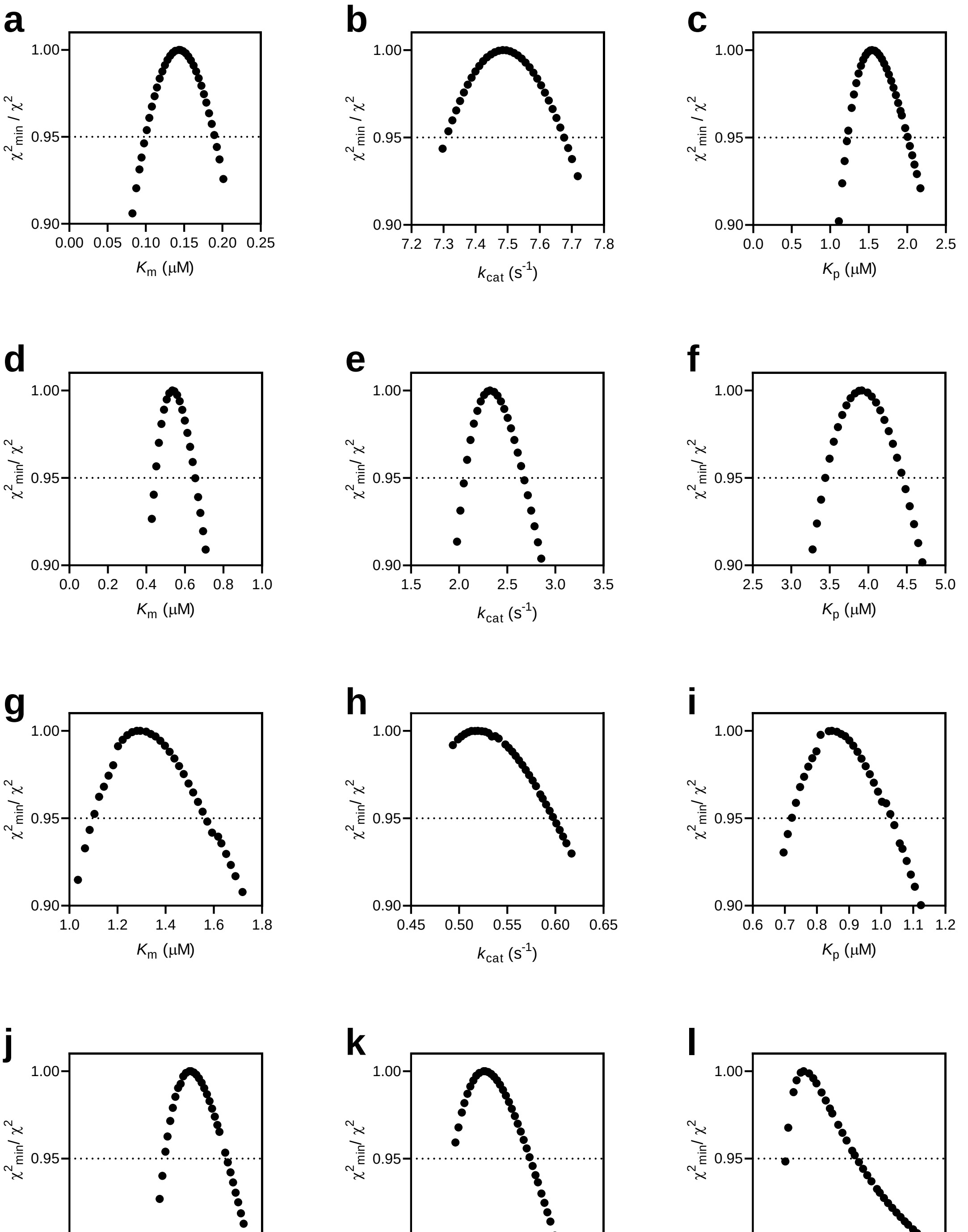


**Supplementary Fig. 13.** Confidence contour analysis of Michaelis constant $K_m$, turnover number $k_{cat}$ and enzyme-product complex dissociation constant $K_p$ obtained as the result of the numerical analysis of kinetic data of NanoLuc catalyzed conversion of FMZ (a, b, c) and CTZ (d, e, f) and NanoLuc-Y94A catalyzed conversion of FMZ (g, h, i) and CTZ (j, k, l), confirming that the determined kinetic parameters are well defined and constrained by the collected kinetic data. The grey dashed lines represent the $\chi^2$ threshold of 0.95.

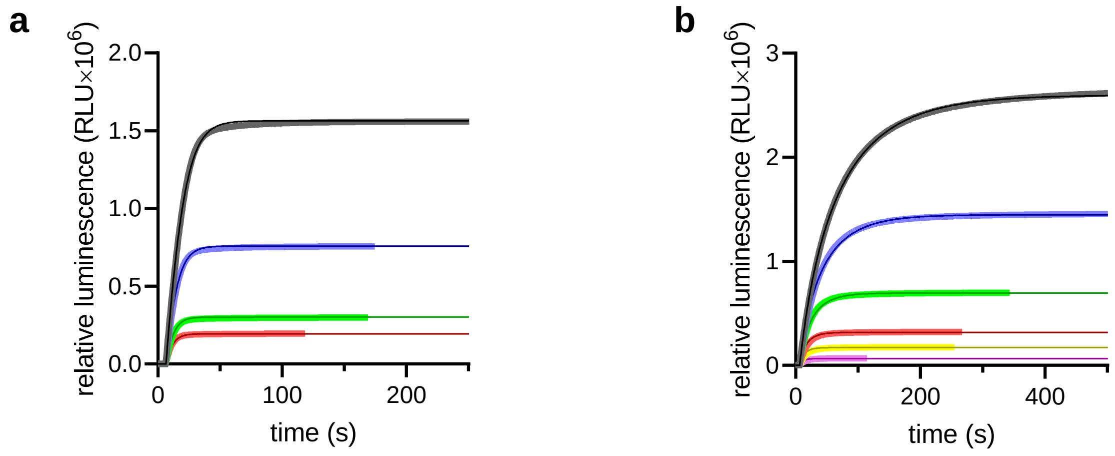


**Supplementary Fig. 14.** Numerical analysis of kinetic parameters of NanoLuc-D9R/K89R catalyzed luciferin conversion. (a) The reaction progress curves corresponding to cumulative luminescence production in time recorded upon mixing 0.01 µM -D9R/H57A/K89R with 0.50 (black), 0.24 (dark blue), 0.10 (green) and 0.06 (red) µM FMZ, with gain of the reader set to 1250. (b) The reaction progress curves obtained upon mixing 0.01 µM NanoLuc-D9R/K89R with 3.63 (black), 2.02 (dark blue), 0.97 (green), 0.44 (red), 0.24 (yellow) and 0.09 (magenta) µM CTZ, with gain of the reader set to 1000. Each trace represents an average of three repetitions. The thicker lines represent the experimental data, thinner lines represent the best fit using reaction model in Scheme 1 (see Methods section Measurement of steady-state kinetic parameters of luciferase reaction).

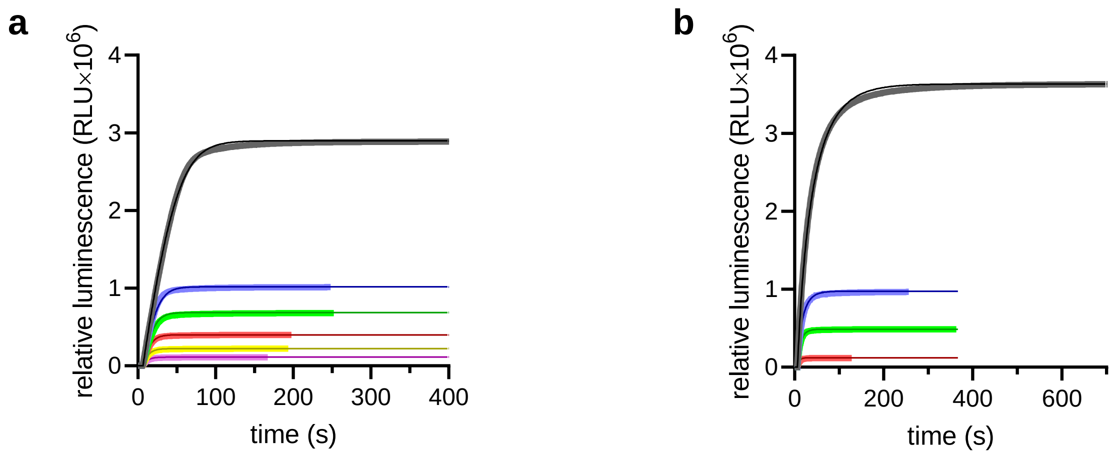


**Supplementary Fig. 15.** Numerical analysis of kinetic parameters of NanoLuc D9R/H57A/K89R-catalyzed luciferin conversion. (a) The reaction progress curves corresponding to cumulative luminescence production in time recorded upon mixing 0.01 µM NanoLuc-D9R/H57A/K89R with 0.98 (black), 0.34 (blue), 0.23 (green), 0.14 (red), 0.08 (yellow) and 0.04 (magenta) µM FMZ, with gain of the reader set to 1250. (b) The reaction progress curves obtained upon mixing 0.01 µM NanoLuc-D9R/H57A/K89R with 3.69 (black), 0.99 (blue), 0.50 (green) and 0.12 (red) µM CTZ, with gain of the reader set to 1000. Each trace represents an average of three repetitions. The thicker lines represent the experimental data, thinner lines represent the best fit using reaction model in Scheme 1 (see Methods section Measurement of steady-state kinetic parameters of luciferase reaction).

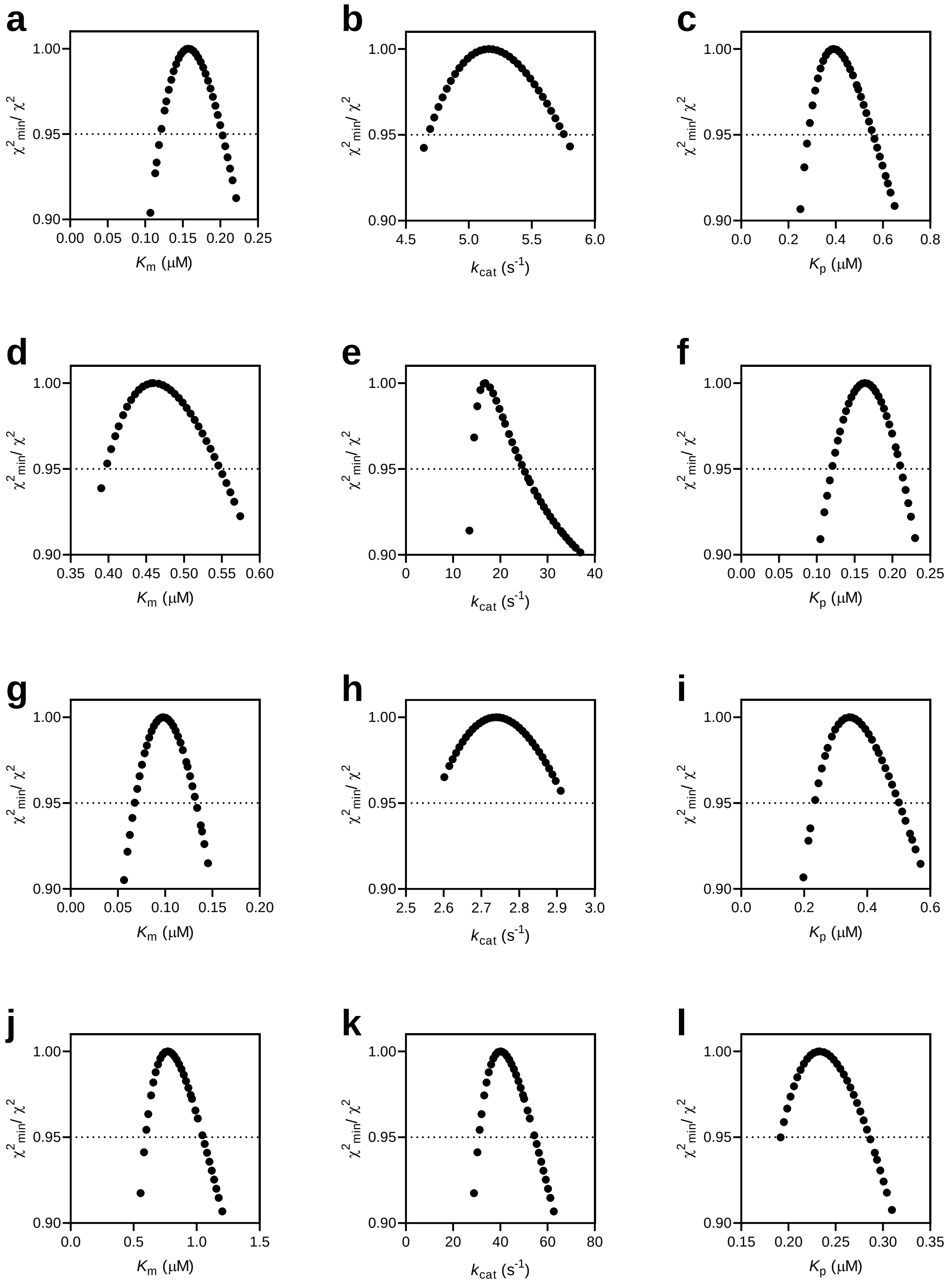


**Supplementary Fig. 16.** Confidence contour analysis of Michaelis constant $K_m$, turnover number $k_{cat}$ and enzyme-product complex dissociation constant $K_p$ obtained as the result of the numerical analysis of kinetic data of NanoLuc D9R/K89R catalyzed conversion of FMZ (a, b, c) and CTZ (d, e, f) and NanoLuc-D9R/H57A/K89R catalyzed conversion of FMZ (g, h, i) and CTZ (j, k, l), confirming that the determined kinetic parameters are well defined and constrained by the collected kinetic data. The grey dashed lines represent the $\chi^2$ threshold of 0.95.

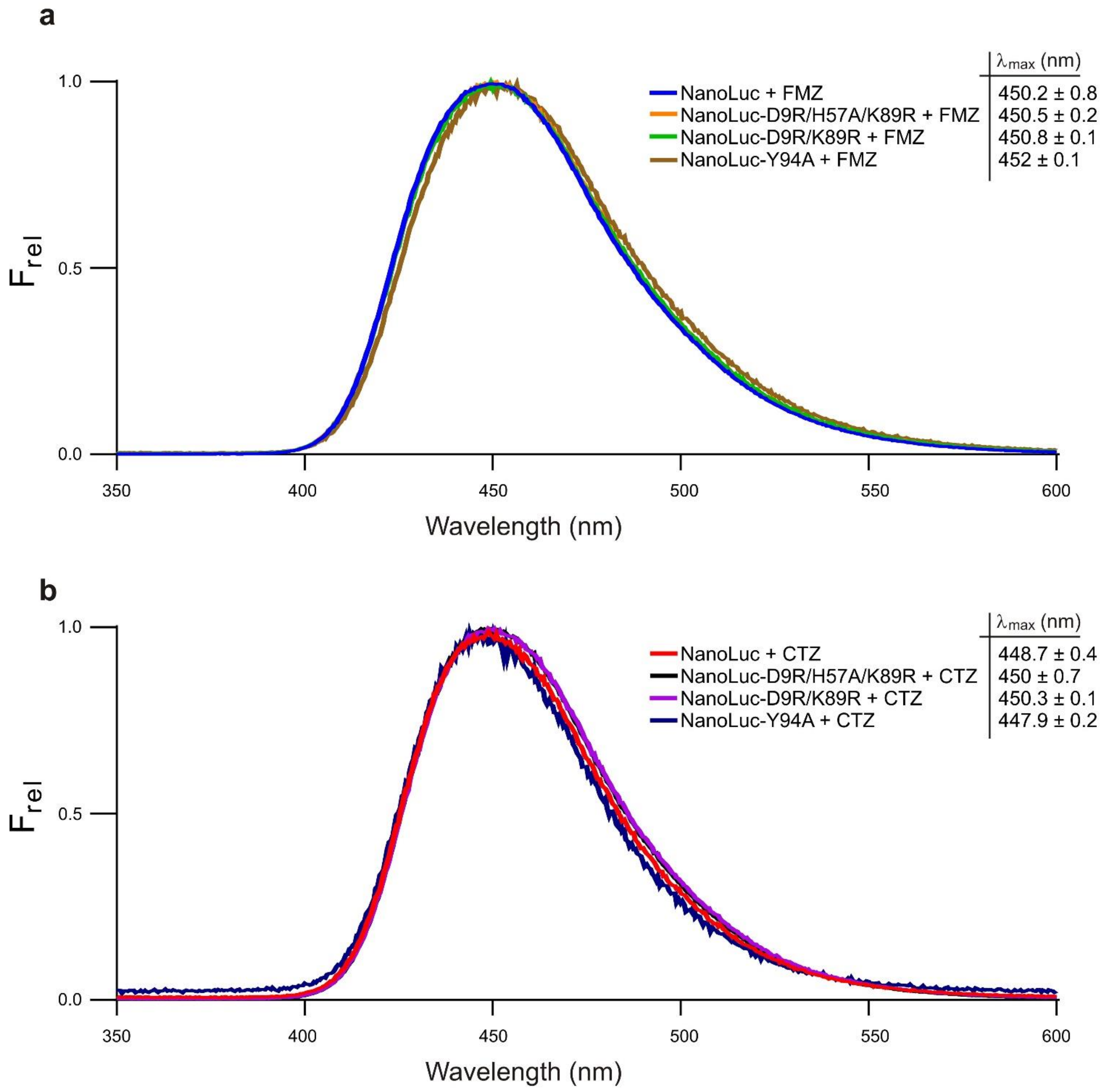


**Supplementary Fig. 17.** Luminescence-emission spectra of NanoLuc and its mutants with FMZ (a) and CTZ (b) luciferins. The values indicate the average of emission maxima calculated from three independent measurements, the error of measurements is represented by the standard deviation (SD).

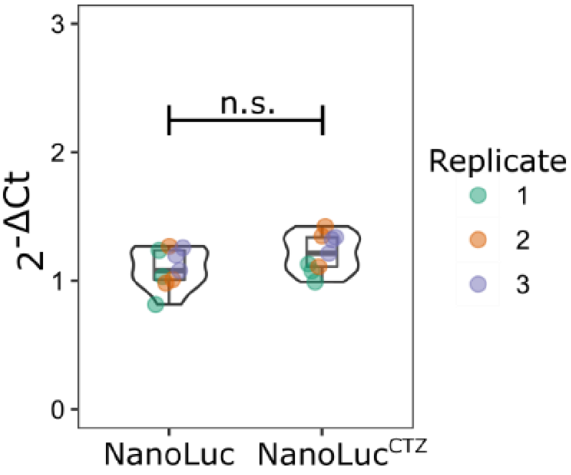


**Supplementary Fig. 18.** Real-time PCR (RT-qPCR) to quantify gene transcripts for NanoLuc and NanoLucCTZ in engineered ARPE-19 cells. Primer sequences used in RT-PCR are shown in **Supplementary Table 8**. Data sets were normalized to the corresponding levels of GAPDH mRNA

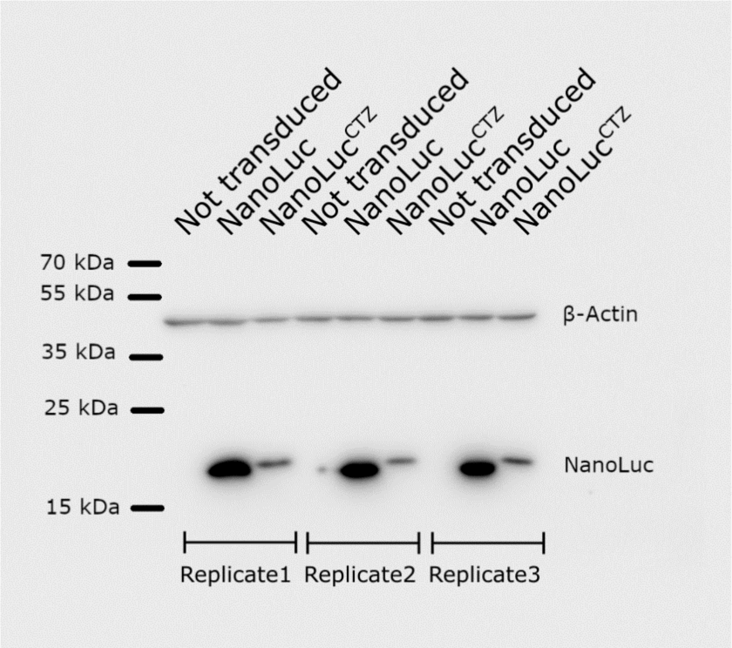


**Supplementary Fig. 19.** Detection of NanoLuc and NanoLuc$^{CTZ}$ expression in ARPE-19 cells, as determined using Western blot analysis. β-Actin represents a loading control. Note that the band intensity for NanoLuc$^{CTZ}$ variant is lower than that of the original NanoLuc. We think that this is because of the primary antibody used that was raised against NanoLuc (N7000, Promega), The NanoLuc$^{CTZ}$ variant contains three mutations on the enzyme surface, which may affect antibody recognition, yielding lower band intensities.

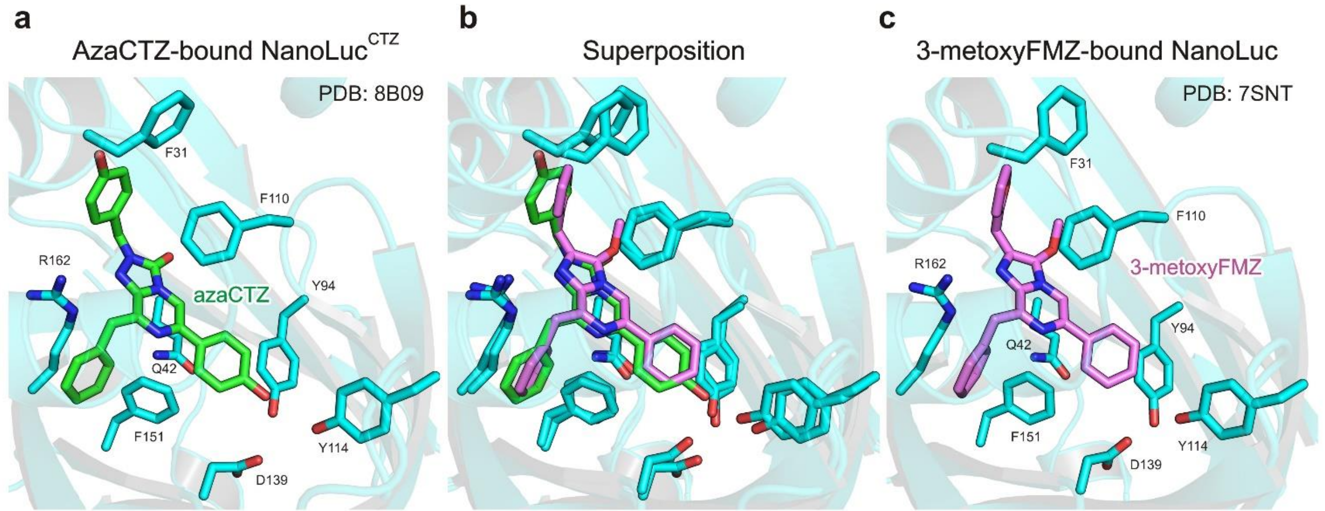


**Supplementary Fig. 20.** Comparison of binding modes of azacoelenterazine (azaCTZ) and 3-methoxy-furimazine (3-metoxyFMZ) when bound in the intra-barrel catalytic site of NanoLuc$^{CTZ}$ (**a**) and NanoLuc (**c**), respectively. Superposition of the two structures is shown in the middle panel (**c**).

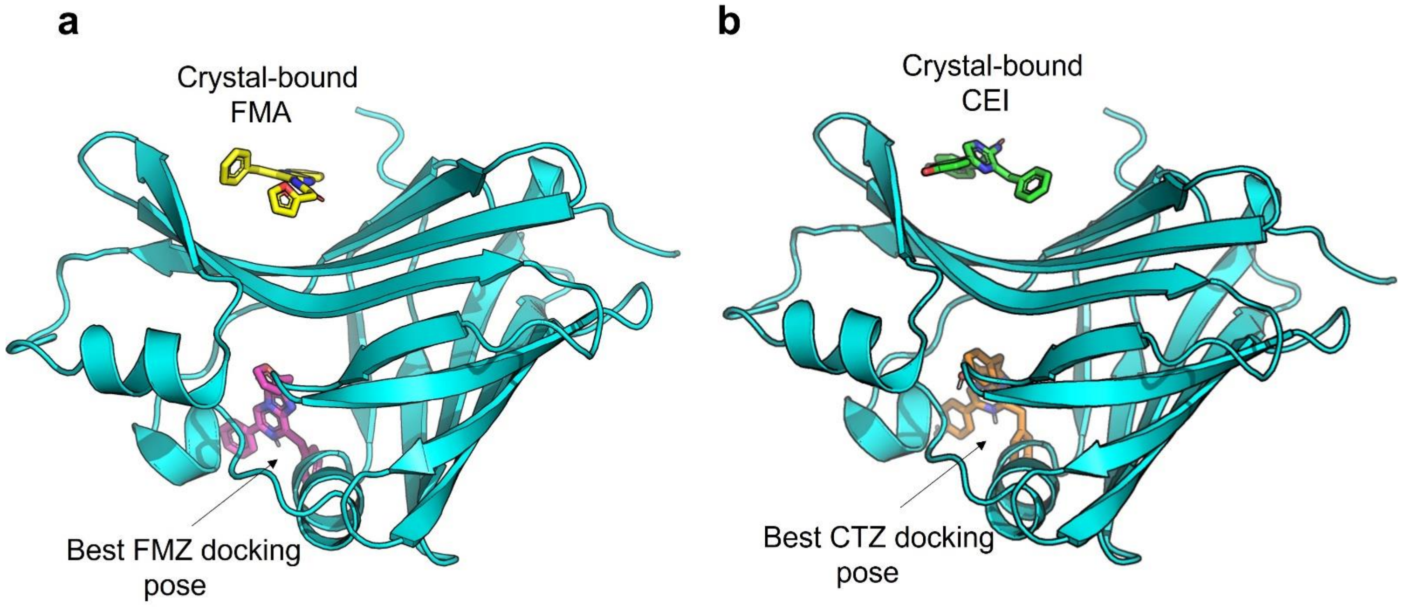


**Supplementary Fig. 21.** The best binding energy poses from blind docking of FMZ (a) and CTZ (b) to monomeric NanoLuc.

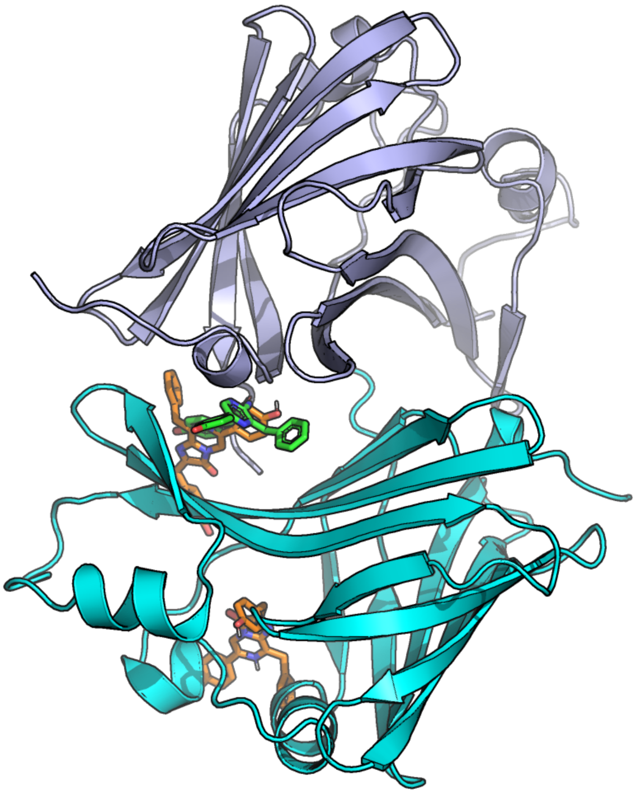

**Supplementary Fig. 22.** Two possible CTZ poses in dimeric NanoLuc. The NanoLuc dimer structure is shown in cartoon representation, with chain A in cyan and chain B in light blue. The crystallographic CEI binding mode is shown as green sticks, while the docked CTZ poses are orange.

**Supplementary Table 3.** Docking of FMZ and CTZ (ligand column) to NanoLuc monomer and dimer (protein form column). The location column specifies the part of the protein to which the pose binds: the entrance vestibule to the central cavity of NanoLuc or the crystallographic binding site in the dimer interface. The energy column describes the predicted binding energy (in kcal/mol) of the pose in that location.

| ligand | protein form | location | energy (kcal/mol) |
|---|---|---|---|
| FMZ | monomer | entrance vestibule | −8.8 |
| | | dimer interface | −6.9 |
| | dimer | entrance vestibule | −8.8 |
| | | dimer interface | −8.5 |
| CTZ | monomer | entrance vestibule | −9.0 |
| | | dimer interface | −7.2 |
| | dimer | entrance vestibule | −9.2 |
| | | dimer interface | −9.3 |

**Supplementary Table 4.** Docking of FMZ to different structures of monomeric NanoLuc using different-sized docking grids: from x = 60 Å, y = 50 Å, z = 46 Å covering the whole protein (blind docking) to gradually smaller grids constraining the docking into the β-barrel of NanoLuc.

| docking grid (Å) | binding energy (kcal/mol) | | | | | |
|---|---|---|---|---|---|---|
| | 8AQ6 | 5IBO | 7MJB | 5B0U | 7VSX | $\text{NanoLuc}^{\text{CTZ}}$ |
| 60 × 50 × 46 | −8.8 | −7.8 | −9.2 | −8.3 | −8.8 | −11.6 |
| 22 × 22 × 22 | 1.9 | −0.4 | −5.5 | −7.5 | −8.8 | −11.6 |
| 20 × 20 × 20 | 7.9 | 6.3 | 0 | −7.5 | −8.6 | −11.6 |
| 18 × 18 × 18 | 17.3 | 7.1 | 0.2 | −6.2 | −8.5 | −11.6 |

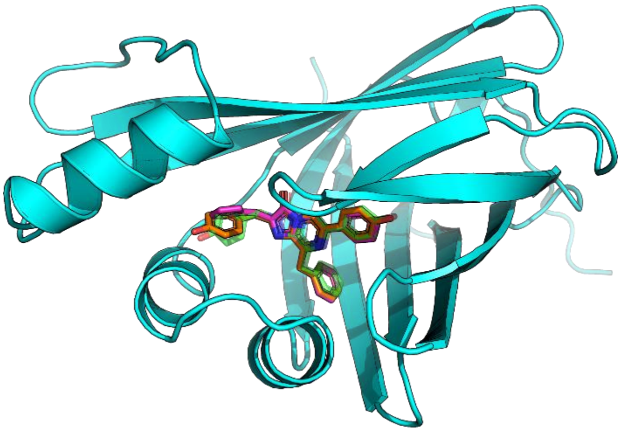

**Supplementary Fig. 23.** The best binding energy poses of FMZ (magenta sticks) and CTZ (orange sticks) from blind docking to NanoLuc$^{CTZ}$ (cyan cartoon) compared to the crystallographic azaCTZ (transparent green sticks).

System: NanoLuc monomer with FMZ

Metric: RMSD of FMZ

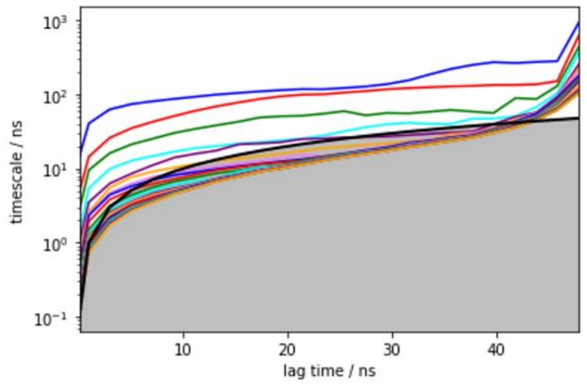


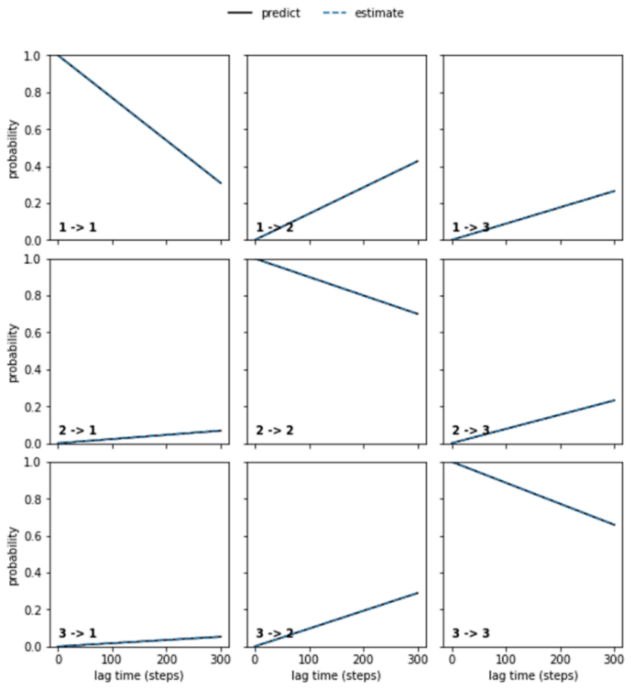


System: NanoLuc dimer with FMZ

Metric: RMSD of FMZ

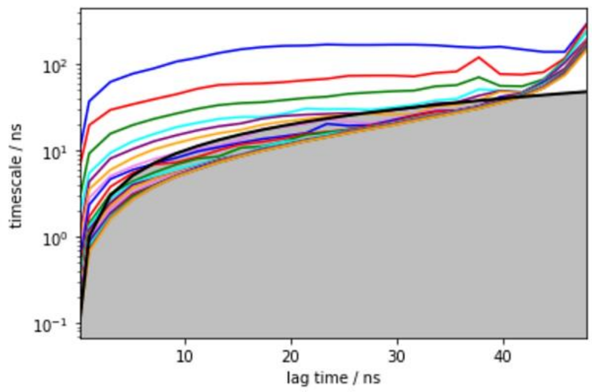


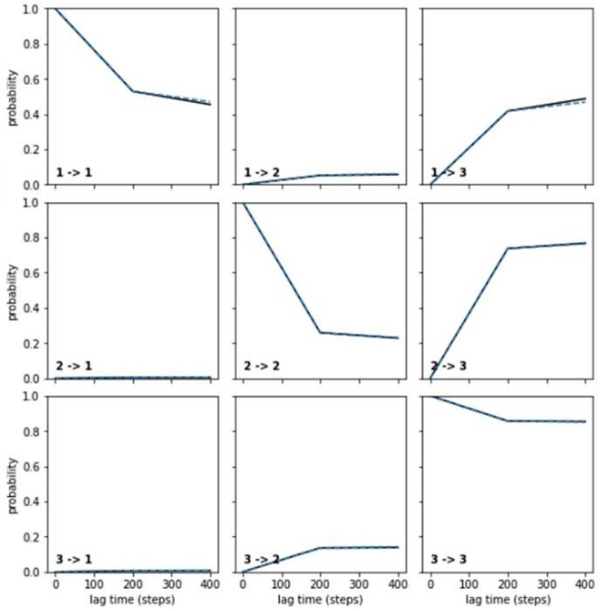


**Supplementary Fig. 24.** Implied timescales and Chapman-Kolmogorov tests for NanoLuc monomer (top row) and NanoLuc dimer with FMZ (bottom row). The metric used for the analysis was the RMSD of FMZ relative to the position of FMA in the crystal structure. The implied timescales of the NanoLuc monomer system (top right) show two transitions, which hints toward three or more macrostates. For NanoLuc dimer, there are three visible transitions in the bottom right plot. Chapman-Kolmogorov tests (left) were done for Markov state models of three macrostates constructed at lag times 30 ns (monomer) and 20 ns (dimer). The selected lag times were the lowest lag times at which the predict (black) and estimate (blue dashed) lines from the Chapman-Kolmogorov test agreed.

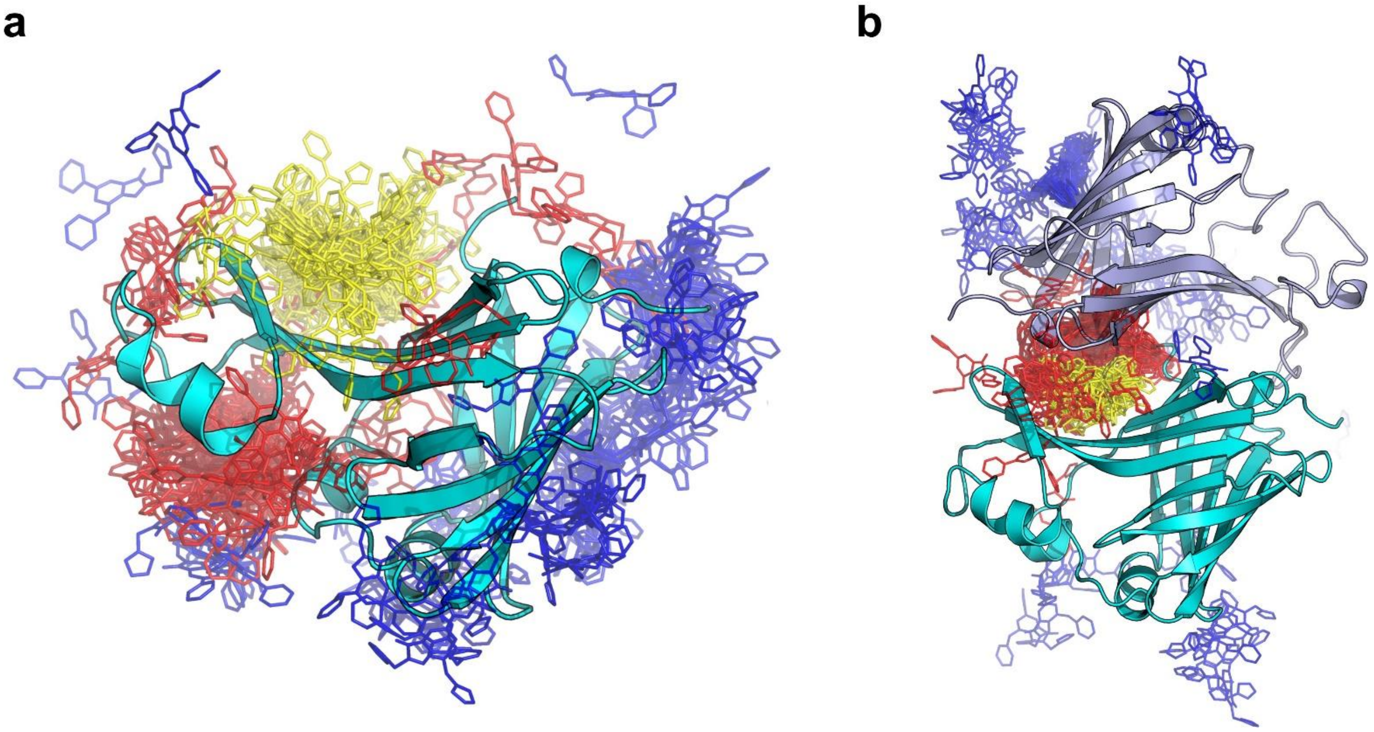


**Supplementary Fig. 25.** Visualization of three macrostates based on the RMSD of FMZ-luciferin (yellow, red, and blue sticks in ascending order of mean RMSD values) from the adaptive sampling of NanoLuc monomer (a) and dimer (b). In the NanoLuc monomer, the yellow macrostate was considered the bound state and the other two as unbound states, while in the dimer, the yellow and red macrostates were considered the bound states and the blue one as the unbound state.

**Supplementary Table 5.** FMZ unbinding kinetics from NanoLuc monomer and dimer. The kinetic parameters were calculated from the Markov state models. For the monomer, the kinetics between the bound (lowest-RMSD state) and the other two states were calculated, while for the dimer, the two lowest-RMSD states were used as the bound states and the third one as the unbound state. The standard deviations were obtained from bootstrapping of random 80 % of the data 100 times.

| | monomer FMZ | dimer FMZ |
|---|---|---|
| $\Delta G^0$* (kcal/mol) | 1.51 ± 0.31 | −1.18 ± 0.27 |
| $k_{on}$* ($M^{-1}\cdot s^{-1}$) | (7.69 ± 3.48) × $10^5$ | (1.79 ± 0.34) × $10^7$ |
| $k_{off}$* ($s^{-1}$) | (9.42 ± 3.81) × $10^6$ | (7.86 ± 9.05) × $10^5$ |
| $k_{off}/k_{on}$ (M) | 12.25 ± 7.43 | 0.04 ± 0.05 |
| $K_D$* (M) | 14.33 ± 18.85 | 0.16 ± 0.08 |

*$\Delta G^0$ – the free energy of the bound state; $k_{on}$ – rate constant of binding; $k_{off}$ – rate constant of unbinding; $k_{off}/k_{on}$ – the ratio of $k_{off}$ to $k_{on}$; $K_D$ – dissociation constant

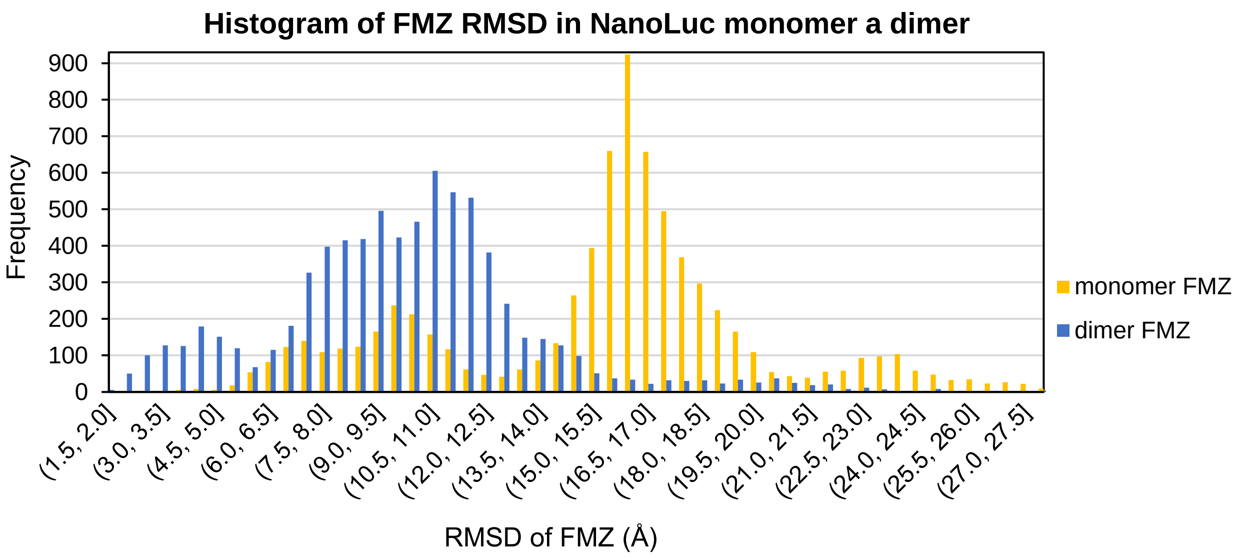


**Supplementary Fig. 26.** Distribution of RMSD values of FMZ in NanoLuc monomer and dimer calculated at every ns of the simulation. FMZ in the dimer (blue columns) exhibits lower RMSD values overall than in the monomer (dark yellow columns), which shows that in the NanoLuc dimer, FMZ prefers staying closer to the crystallographic binding site.

System: NanoLuc dimer with FMZ

Metric: RMSD of protein Cα

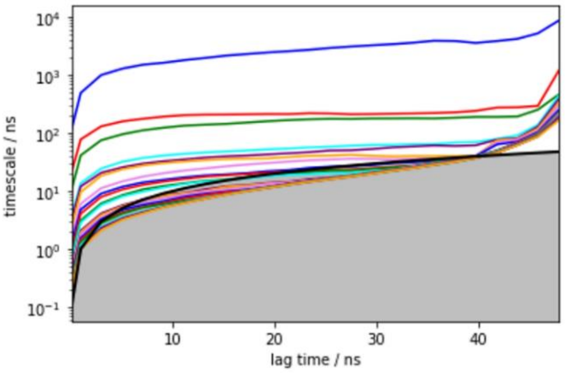


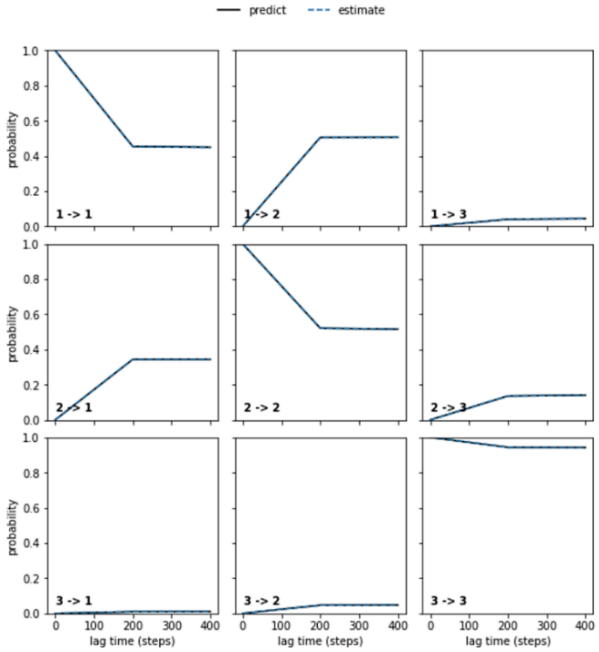


System: NanoLuc dimer alone

Metric: RMSD of protein Cα

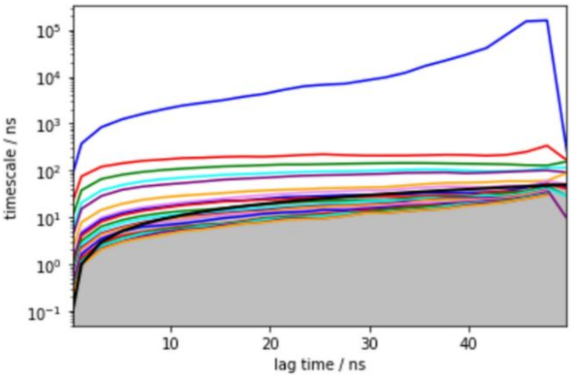


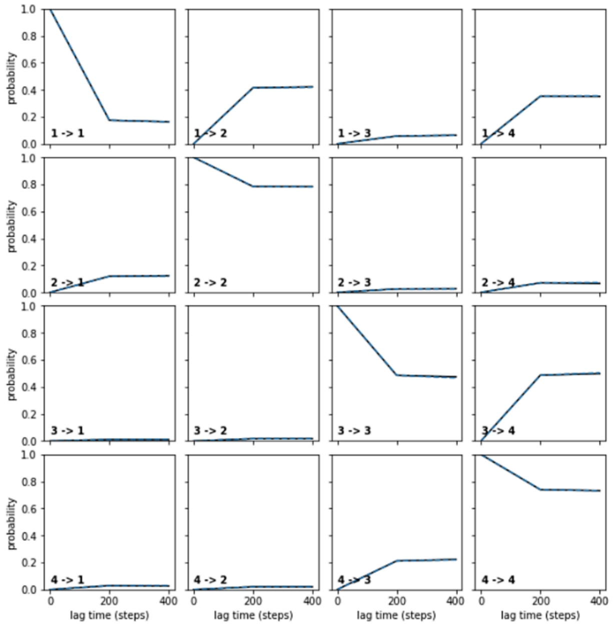


**Supplementary Fig. 27.** Implied timescales and Chapman-Kolmogorov tests of NanoLuc dimer in complex with FMZ (top row) and NanoLuc dimer alone (bottom row). The metric used for the analysis was the RMSD of the protein Cα atoms. The implied timescales of the NanoLuc dimer with FMZ system (top right) show two transitions, which hints toward three or more macrostates. For NanoLuc dimer alone, there is only one clear transition in the bottom right plot. Chapman-Kolmogorov tests (on the left) were done for Markov state models constructed at a lag time of 20 ns (the lowest lag time at which the black predict and blue dashed estimate lines from the Chapman-Kolmogorov test agreed). Three macrostates were constructed from the dimer with FMZ simulation (top right) and four macrostates from the dimer alone (bottom right).

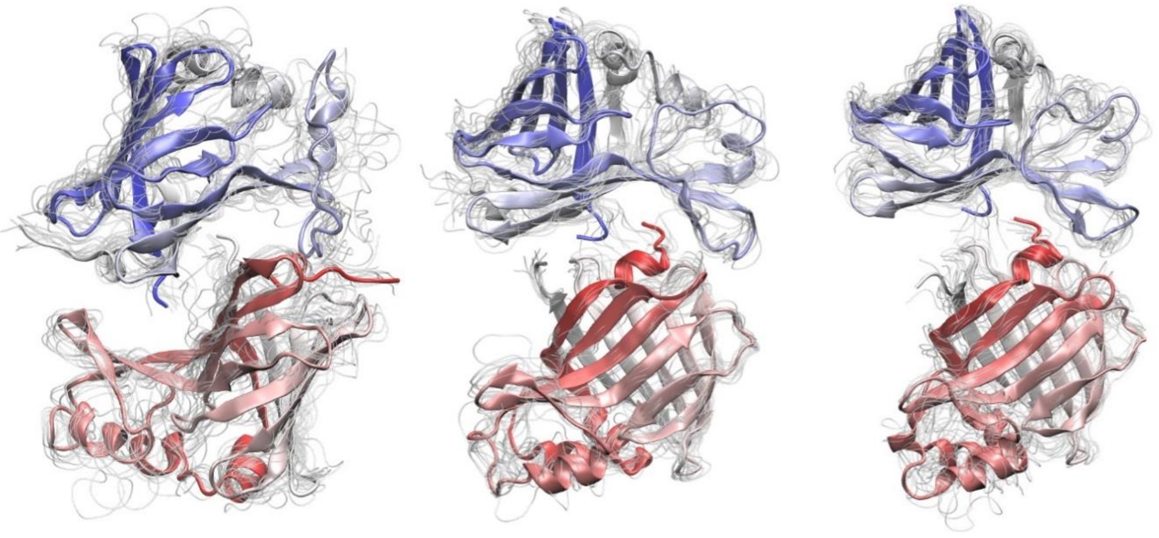

**Supplementary Fig. 28.** Three macrostates from simulations with NanoLuc dimer in complex with FMZ. Each macrostate is visualized with a representative structure shown as an index-colored cartoon, and to show the variance, every tenth snapshot was visualized as a thin transparent light gray tube. The most crystal-like state (associated state) is on the left with a mean RMSD of 4.3 Å, while the partially dissociated macrostates in the middle and on the right exhibit a mean RMSD of 10.9 and 17.1 Å, respectively.

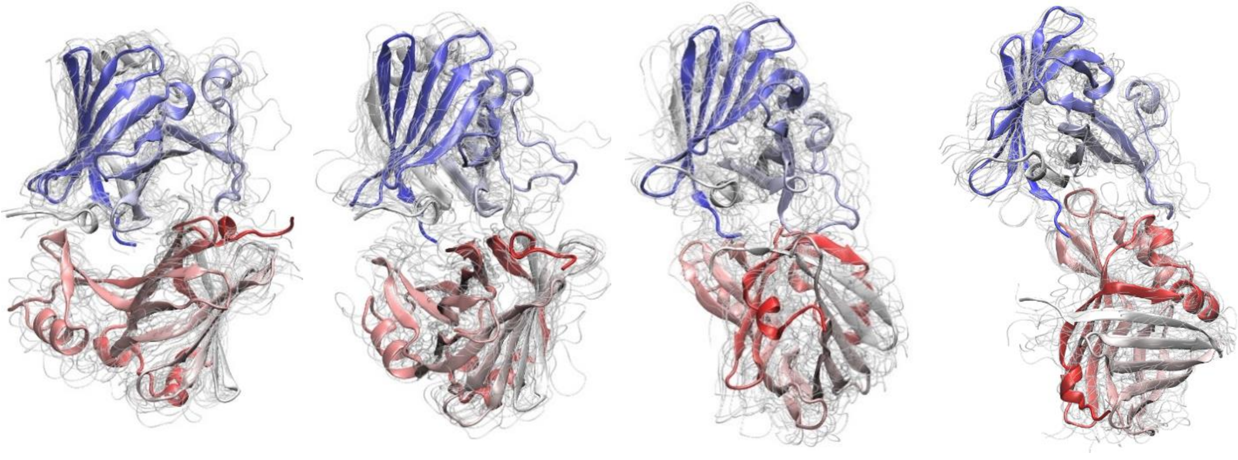

**Supplementary Fig. 29.** Four macrostates from NanoLuc dimer simulation without FMZ. Each macrostate is visualized with a representative structure shown as an index-colored cartoon and every tenth snapshot shown as a thin transparent light gray tube. The most left macrostate is the most associated one with a mean RMSD of 4.9 Å, which is comparable with the lowest-RMSD macrostate from **Supplementary** Chyba! Nenalezen zdroj odkazů. (macrostates of NanoLuc dimer with FMZ). The remaining macrostates show gradual dissociation with mean RMSD values of 9.7, 15.0, and 19.8 Å, from left to right.

**Supplementary Table 6.** Dimer dissociation kinetics of NanoLuc dimer with and without FMZ. The kinetic parameters were calculated from the Markov state models. The kinetics were calculated between the macrostates with the lowest and highest mean RMSD. The standard deviations were obtained from bootstrapping of random 80 % of the data 100 times.

| | dimer FMZ | dimer |
|---|---|---|
| $\Delta G^0$* (kcal/mol) | −0.96 ± 0.49 | 0.02 ± 0.57 |
| $k_{as}$* ($M^{-1}\cdot s^{-1}$) | (3.58 ± 1.22) × $10^5$ | (5.03 ± 1.87) × $10^4$ |
| $k_{dis}$* ($s^{-1}$) | (1.03 ± 0.51) × $10^5$ | (1.19 ± 0.43) × $10^5$ |
| $k_{dis}/k_{as}$* (M) | 0.29 ± 0.17 | 2.35 ± 1.23 |
| $K_D$* (M) | 0.28 ± 0.22 | 1.50 ± 1.39 |

*$\Delta G^0$ – the free energy of the associated state; $k_{as}$ – rate constant of association; $k_{dis}$ – rate constant of dissociation; $k_{dis}/k_{as}$ – the ratio of $k_{dis}$ to $k_{as}$; $K_D$ – dissociation constant

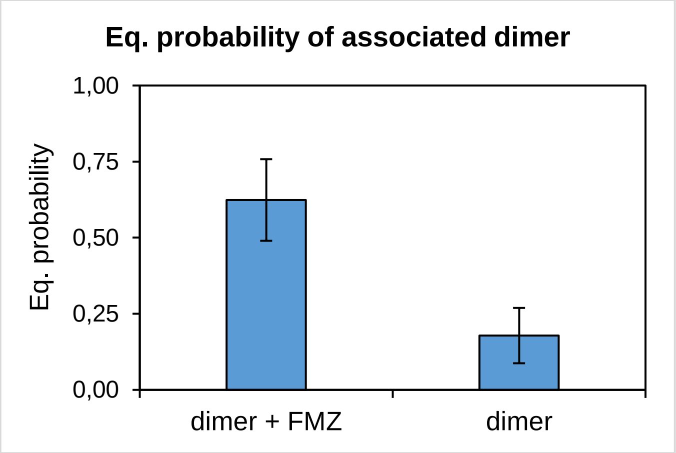


**Supplementary Fig. 30.** Equilibrium probability of associated dimer macrostates from simulation of dimeric NanoLuc with (left column) and without FMZ (right column). The associated macrostate corresponds to the lowest-RMSD macrostate. The error bars show the standard deviation from bootstrapping calculation with a random 80 % of the data repeated 100 times. This plot shows that the associated dimer is significantly more probable when FMZ is present in the simulation.

**Supplementary Table 7.** List of used primers in PCR-based mutagenesis experiments.

| Mutation | Mutagenic primers (nucleotide sequence 5'-3') |
|---|---|
| R43A | GTTAGCGTTACCCCGATTCAGGCGATTGTTCTGAGCGGTGAAAAT |
| R11E | GAAGATTTTGTTGGTGATTGGGAGCAGACCGCAGGTTATAATCTG |
| D55A | GGTGAAAATGGCCTGAAAATTGCGATTCATGTGATCATCCCGTAT |
| H57A | GAAAATGGCCTGAAAATTGATATTGCGGTGATCATCCCGTATGAAGGTCTG |
| Y81A | GAAAAAATCTTCAAAGTTGTGGCGCCGGTGGATGACCACCATTTT |
| K89A | GGTGCCATAATGCAGAATCACCGCAAAATGGTGGTCATCCACCGG |
| K89E | CAGGGTGCCATAATGCAGAATCACCTCAAAATGGTGGTCATCCACCGGATA |
| Y94A | ACCATCAATAACCAGGGTGCCCGCATGCAGAATCACTTTAAAATG |
| R166A | GAATTCGGATCCTTATGCCAGAATCGCTTCACACAGACGCCAACCGGTAAC |
| D9R | TTTACCCTGGAAGATTTTGTTGGTCGCTGGCGTCAGACCGCAGGTTATAAT |
| H57Y | GAAAATGGCCTGAAAATTGATATTTACGTGATCATCCCGTATGAAGGTCTG |
| K89R | CAGGGTGCCATAATGCAGAATCACCCTAAAATGGTGGTCATCCACCGGATA |
| R166Q | GAATTCGGATCCTTATGCCAGAATCTGTTCACACAGACGCCAACCGGTAAC |
| del I167-A169 | ACGGAGCTCGAATTCGGATCCTTAACGTTCACACAGACGCCAACCGGT |
| ext A170-A171 | TCGACGGAGCTCGAATTCGGATCCTTACGCCGCTGCCAGAATACGTTCACACAGACG |
| R162A | ACAATTAATGGTGTTACCGGTTGGGCGCTGTGTGAACGTATTCTGGCATAA |
| Y109A | GGTGTGACCCCGAATATGATTGATGCGTTCGGTCGTCCGTATGAGGGTATT |
| D139A | CTGTGGAACGGCAATAAAATCATTGCGGAACGTCTGATTAATCCGGATGGT |
| Universal primers (nucleotide sequence 5'-3') | |
| forward | TAATACGACTCACTATAGGG |
| reverse | GCTAGTTATTGCTCAGCGG |

**Supplementary Table 8.** List of used primers in RT-PCR experiments.

| Mutation | Mutagenic primers (nucleotide sequence 5'-3') |
|---|---|
| GAPDH-F | TGCACCACCAACTGCTTAGC |
| GAPDH-R | GGCATGGACTGTGGTCATGAG |
| NanoLuc-F | GCTATAGCTAGCACCATGCATCATCATCACCATCACAGCG |
| NanoLuc-R | GGTGCCATAATGCAGAATCACCGCAAAATGGTGGTCATCCACCGG |

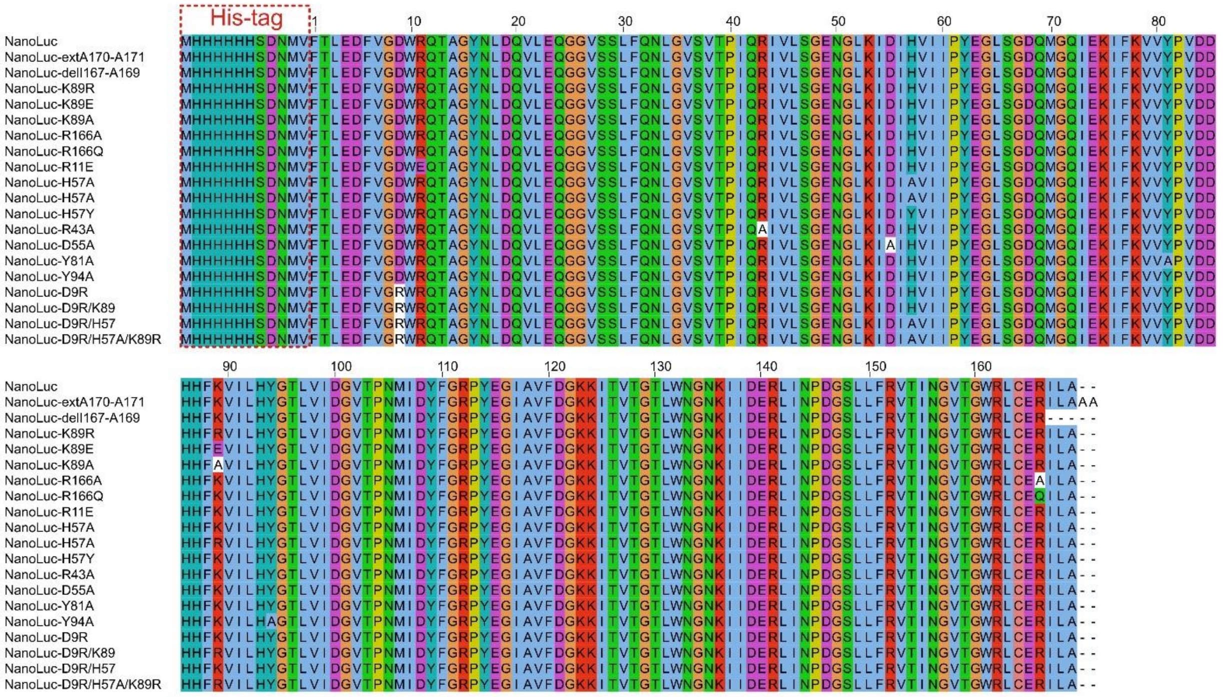

**Supplementary Fig. 31.** Sequence alignment of all cloned and analyzed NanoLuc mutants in this work.

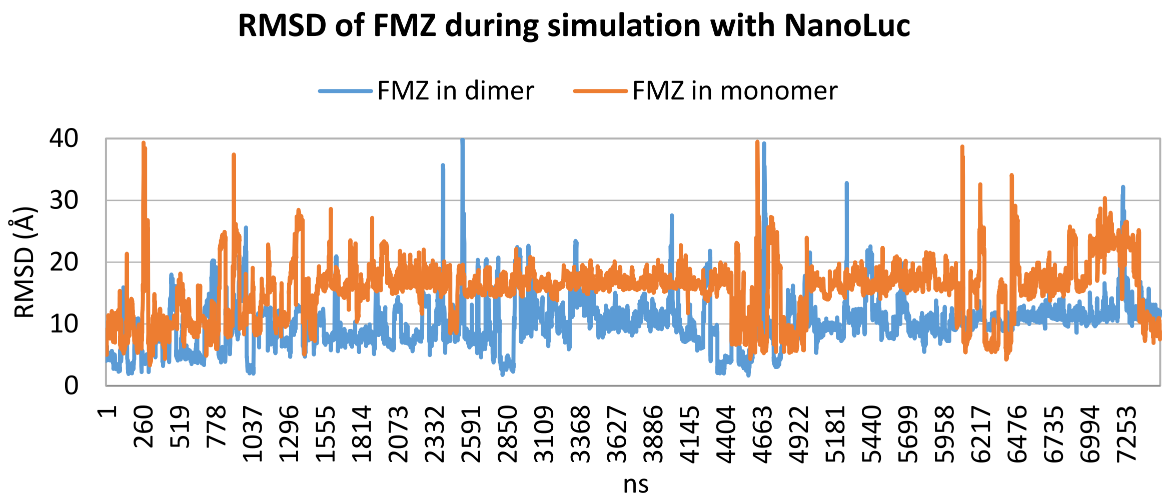


**Supplementary Fig. 32.** Time-course analysis of the RMSD of furimazine from its binding pose in the allosteric site. The RMSD of FMZ simulated with dimeric NanoLuc is marked by the blue line, while the orange line is from simulation with NanoLuc monomer.

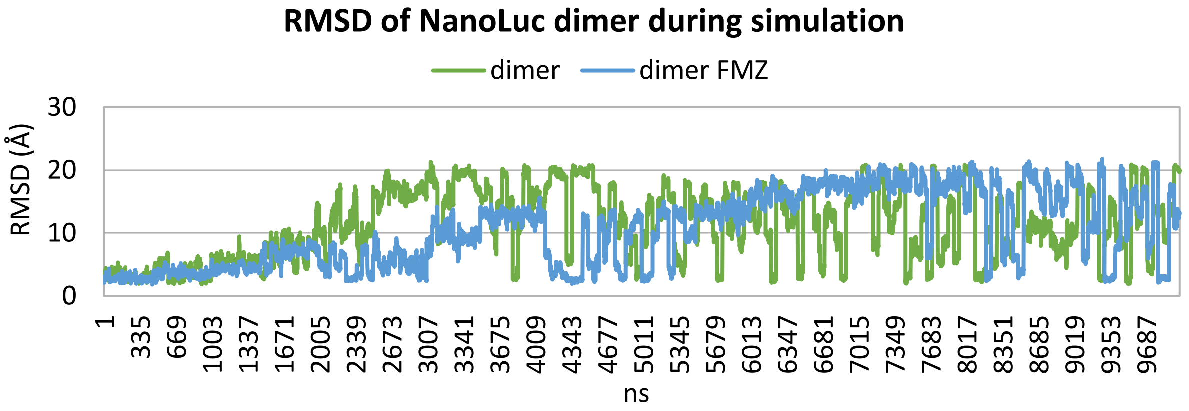


**Supplementary Fig. 33.** Time-course analysis of the RMSD of NanoLuc dimer from its associated state, as observed in the crystal structure. The RMSD of NanoLuc dimer simulated with furimazine is shown by the blue line, while the green line is from simulation without ligand.

**Supplementary Schemes 1 and 2**

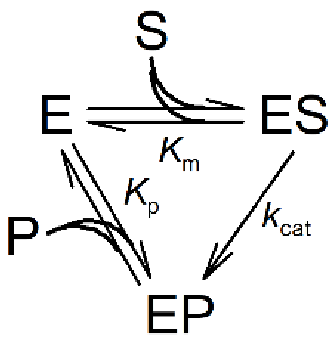


Supplementary Scheme 1

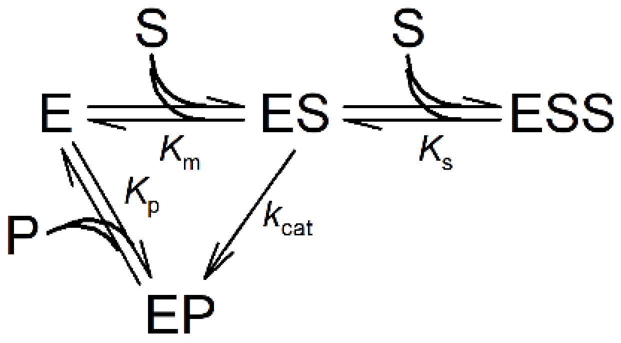


Supplementary Scheme 2

**Supplementary Note 1. Synthesis of azafurimazine.**

For azafurimazine (azaFMZ) synthesis, $^{1}$H NMR and $^{13}$C NMR spectra were recorded on a Bruker Avance 400 spectrometer at 400 MHz and 100 MHz, respectively. Shifts (δ) are given in ppm with respect to the TMS signal and cross-coupling constants (J) are given in Hertz. Column chromatography were performed either on Merck silica gel 60 (0.035 - 0.070 mm) or neutral alumina containing 1.5% of added water using a solvent pump and an automated collecting system driven by a UV detector set to 254 nm unless required otherwise. Sample deposition was carried out by absorption of the mixture to be purified on a small amount of the solid phase followed by its deposition of the top of the column. The low-resolution mass spectra were obtained on an Agilent 1200 series LC/MSD system using an Agilent Jet-Stream atmospheric electrospray ionization system and the high-resolution mass spectra (HRMS) were obtained using a Waters Micromass Q-Tof with an electrospray ion source.

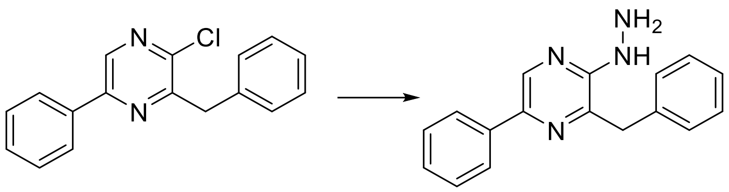

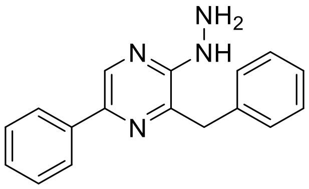

3-benzyl-2-hydrazinyl-5-phenylpyrazine: In a 20 mL sealable Biotage vial, 3-benzyl-2-chloro-5-phenylpyrazine[41] (2.0 g, 7.1 mmol) and hydrazine hydrate (1.38 mL, 28.5 mmol) were dispersed in *n*-butanol (12 mL). The vial was sealed and heated in a microwave oven at 170 °C for 8 h. The resulting mixture was dispersed in distilled water (200 mL) for 15 minutes at room temperature. The precipitated was then filtered, washed with distilled water, cyclohexane and dried under vacuum at 55°C to give the hydrazine as a yellow solid (3.71 g, 93%). $^{1}$H (DMSO-$d_6$) $\delta$ 8.55 (s, 1H), 8.07 (s, 1H), 7.95-7.92 (m, 2 H), 7.44-7.17 (m, 9H), 4.30 (s (br), 2H), 4.12 (s, 2H). $^{13}$C (DMSO-$d_6$) $\delta$ 153.7, 141.3, 138.8, 138.3, 137.5, 136.4, 129.5, 129.1, 128.6, 127.9, 127.2, 126.7, 125.3. HRMS (m/z): $[M+H]^+$ calcd for $C_{17}H_{16}N_4$: 277.1453, found: 277.1458.

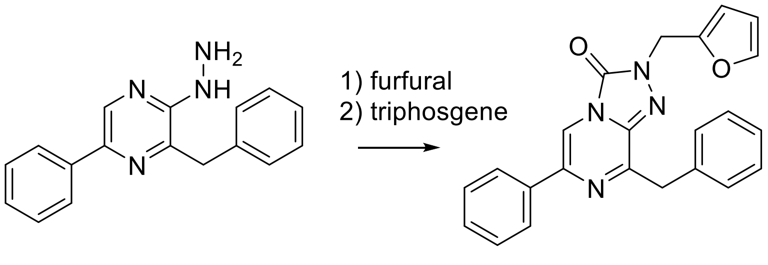

8-benzyl-2-(furan-2-ylmethyl)-6-phenyl-[1,2,4]triazolo[4,3-a]pyrazin-3(2H)-one (azafurimazine). First step: in a 250 mL round bottomed flask, 3-benzyl-2-hydrazinyl-5-phenylpyrazine (0.4 g, 1.45 mmol) and furan-2-carbaldehyde (0.12 mL, 1.45 mmol) were dispersed in acetic acid (4 mL). The mixture was stirred at room temperature during 2 minutes. The resulting solid was re-dissolved in dichloromethane (40 mL) and cyanoborohydride (0.18 g, 2.9 mmol) was added. This was stirred at room temperature for 1 h. The solution was then dispersed in water and ethyl acetate, neutralized with 1 N NaOH (1 equivalent in regard with the acetic acid added). This was extracted with ethyl acetate thrice, the organic layer was washed with a saturated solution of sodium hydrogenocarbonate, distilled water, brine and dried over $MgSO_4$. The solvent was removed under vacuum, to give the crude hydrazine (0.43 g) as a brown oil which was considered pure. Second step: under an inert atmosphere, this oil was dissolved in dry tetrahydrofuran (20 mL, dried over 4 Å molecular sieves) and solid triphosgene (0.12 g, 0.40 mmol) was added before stirring at room temperature for 40 minutes. The resulting mixture was diluted with water, extracted twice with ethyl acetate and the organic layer washed with water, brine, dried over magnesium sulfate and concentrated under vacuum to give a solid residue. A chromatography over silica gel (cyclohexane - ethyl acetate 5/1) gave a fraction containing pure azafurimazine as a yellow solid (0.37 g, 80%). $^{1}H$ ($CDCl_3$) $\delta$ 7.91-7.86 (m, 3H), 7.53-7.24 (m, 9H), 6.48 (dd, $J$ = 3.3, 0.6 Hz, 1H), 6.39 (dd, $J$ = 3.2, 1.9 Hz, 1H), 5.25 (s, 2H), 4.37 (s, 2H). $^{13}C$ ($CDCl_3$) $\delta$ 154.0, 148.5, 148.4, 143.0, 136.5, 136.0, 135.7, 135.5, 129.7, 128.9, 128.7, 128.5, 126.9, 125.7, 110.6, 109.5, 109.3, 43.0, 39.5. HRMS (m/z): $[M+H]^+$ calcd for $C_{23}H_{18}N_4O_2$: 383.1508, found: 383.1494.
